# Giant DNA viruses encode a hallmark translation initiation complex of eukaryotic life

**DOI:** 10.1101/2025.09.30.678621

**Authors:** J. Maximilian Fels, Aidan B. Hill, Richard Han, Jasmine M. Garcia, Hugo Bisio, Chantal Abergel, Philip J. Kranzusch, Amy S.Y. Lee

## Abstract

In contrast to living organisms, viruses were long thought to lack protein synthesis machinery and instead depend on host factors to translate viral transcripts. Here, we discover that giant DNA viruses encode a distinct and functional IF4F translation initiation complex to drive protein synthesis, thereby blurring the line between cellular and acellular biology. During infection, eukaryotic IF4F on host ribosomes is replaced by an essential viral IF4F that regulates viral translation, virion formation, and replication plasticity during altered host states. Structural dissection of viral IF4F reveals that the mRNA cap-binding subunit mediates exclusive interactions with viral mRNAs, constituting a molecular switch from translating host to viral proteins. Thus, our study establishes that viruses express a eukaryotic translation initiation complex for protein synthesis, illuminating a series of evolutionary innovations to a core process of life.

## Introduction

The capacity to produce proteins at the appropriate level, timing, and location is vital for all cellular functions across the kingdoms of life. Indeed, genes related to protein synthesis make up the majority of the universal gene set of life, demonstrating the ancient and conserved role of translation in cellular organisms^1,2^. Building upon these primordial origins, the increased complexity of modern organisms has been supported by the evolution of mRNA translation mechanisms that tune the proteome for specialized cellular functions. For instance, in eukaryotes, defining adaptations to protein synthesis include m^7^G-capped mRNAs and a heterotrimeric cap-binding complex named eIF4F, which regulates the rate-limiting step of translation through the coordinated functions of the cap-binding protein eIF4E, the scaffolding protein eIF4G, and the DEAD-box helicase eIF4A^3–6^. In contrast to living organisms, viruses cannot replicate independently and rely on a host cell to perform many of the biological processes required to reproduce^7^. Although viruses encode proteins involved in DNA replication and transcription, the dogma is that all viruses share a universal dependence on the host cell translation machinery for viral protein synthesis. This principle was challenged by the discovery of nucleocytoplasmic large DNA viruses of the *Megaviricetes* class^8,9^. These viruses infect a broad range of protist and metazoan hosts and are termed “giant DNA viruses” due to possessing genome and particle sizes approaching those of bacteria. Astonishingly, giant DNA viruses collectively encode hundreds of genes with homology to tRNAs, aminoacyl-tRNA synthetases (aaRS), and even putative eukaryotic translation factors^10,11^. Given the low sequence similarity between these proteins and eukaryotic translation factors, as well as the absence of such hallmark proteins of cellular life in other viruses, it remains unclear why giant viruses encode translation factors and whether these factors represent features related to the emergence of eukaryotic protein synthesis. Here, we discover that giant DNA viruses encode an IF4F mRNA cap-binding complex, characteristic of eukaryotic protein synthesis, that controls translation during infection. Viral IF4F coordinates synthesis of structural proteins during late stages of infection, and this transcript-specific regulation occurs through evolutionary innovations to viral IF4F-mediated m^7^G cap recognition. We further show that the viral IF4F complex allows for unparalleled replication efficiency under stress conditions, suggesting that viral expression of eukaryotic-like translation machinery drives a robust evolutionary advantage.

## Results

### Discovery of a viral translation initiation complex

To define the boundaries of viral and cellular functions in protein synthesis and to identify novel forms of translation regulation, we leveraged the prototypical giant DNA virus mimivirus (*Acanthamoeba polyphaga mimivirus*, APMV, **Fig. 1A**) as a model system. APMV infects single cell protists of the Acanthamoeba genus, and we hypothesized that a bona-fide translation factor encoded by APMV would interact with host ribosomes during infection. We examined translation complex composition from mock-infected and APMV-infected cells 8 hours post-infection (hpi) by polysome profiling (**Fig. 1B, Fig. S1D**). As expected, mass spectrometry of the 40S fraction identified host proteins which were enriched in translation and metabolism-related gene ontologies^12^ (**Fig. 1C, Fig. S2F–G**). Among the viral proteins, two with sequence homology to eIF4A (R458) and eIF4E (L496) co-sedimented with 40S small ribosomal subunits during infection. This ribosome association suggests a role as functional translation factors, and we therefore designate these proteins viral IF4A (vIF4A) and viral IF4E (vIF4E) (**Fig. 1C, Fig. S1A–C**). Mass spectrometry analyses also revealed 54 ribosome-associated viral proteins that lack detectable sequence homology to any characterized protein (**Table S1**). Since the complex origins and high mutation rates of viral lineages can reduce the ability to derive function from sequence, we leveraged a structural homology search to de-orphan these proteins. Using AlphaFold2 and Foldseek analysis followed by manually filtering for candidates with translation-related functions, we made the surprising discovery that viral protein R255 is a predicted structural homolog of the middle domain of eIF4G (MIF4G). To validate this finding, we determined a 1.5 Å crystal structure of R255 which revealed a fold consisting of tandem α-helical HEAT repeats (**Fig. 1D**). Structural comparison confirmed strong homology between R255 and MIF4G (TM score = 0.64), and we therefore designate this protein viral IF4G (vIF4G) (**Fig. 1E**). Homologs of vIF4G are also present in giant viruses of the *Klosneuvirinae* subfamily, but phylogenetic analysis revealed that vIF4G from APMV and other members of the *Megamimivirinae* subfamily are part of a separate and highly divergent clade (**Fig. S2A**).

**Figure 1.**
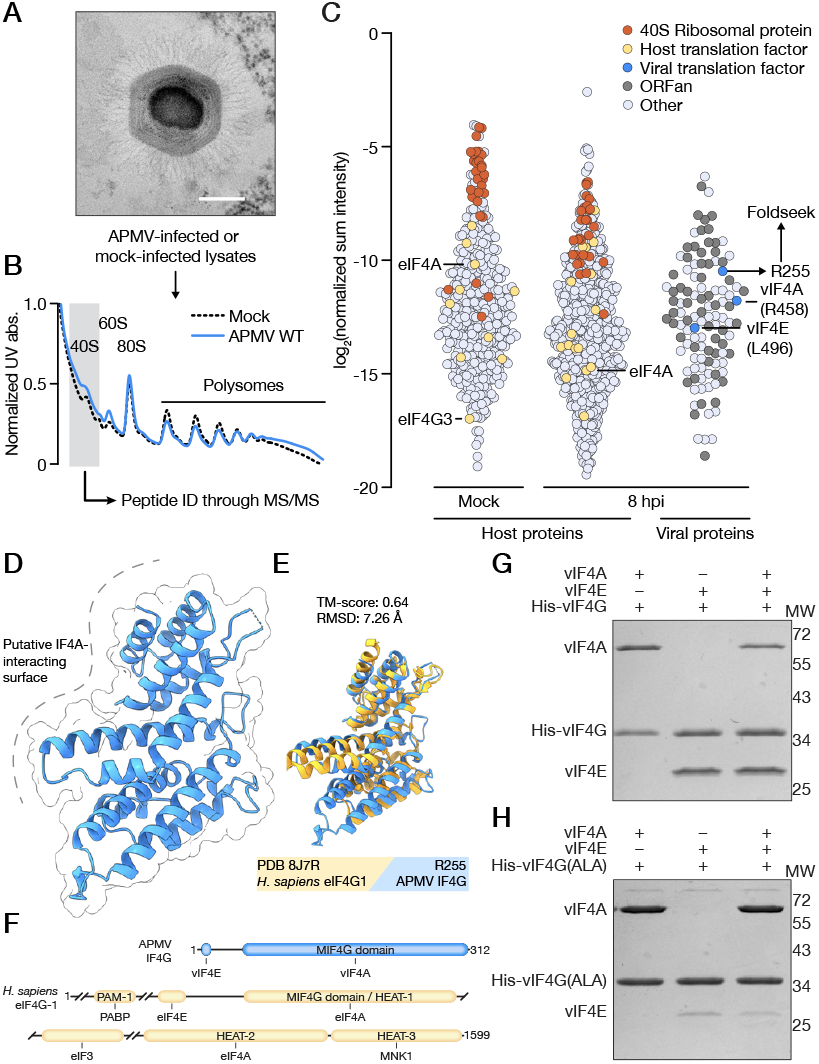
Discovery of a viral translation initiation complex. **A**. TEM micrograph of a mimivirus particle and a schematic of the isolation of 40S ribosomes from infected cells. Bar = 200 nm. **B**. Polysome profiles of lysates from mock-infected and APMV WT-infected cells. **C**. Normalized abundances of host and viral proteins from 40S ribosomal fractions. Each circle represents a single protein, colored accordingly: 40S ribosomal proteins in orange, host translation factors in yellow, viral translation factors in blue, viral ORFan proteins in dark grey, all other proteins, host and viral, in light grey. **D**. X-ray crystal structure of APMV vIF4G (R255) at 1.46 Å resolution (PDB 9E5A). **E**. Structural comparison of APMV vIF4G and the *Hs*. eIF4G1 MIF4G domain (PDB 8J7R) generated using Foldseek. **F**. Domain organization of APMV vIF4G and *Hs*. eIF4G1. **G**. vIF4F complex formation mediated by His-vIF4G visualized by Coomassie-stained SDS-PAGE gel. **H**. Dissection of the interaction between vIF4E and His-vIF4G(ALA), in which the YX_3_II motif has been mutated to alanine, visualized by Coomassie-stained SDS-PAGE gel. Results in G and H are representative of two independent experiments.

eIF4G utilizes a complex multi-domain architecture to act as a scaffolding protein requisite for formation of eIF4F and recruitment of the mRNA-bound complex to the 40S ribosome (**Fig. 1F**). We therefore asked if vIF4G, despite being highly truncated compared to eIF4G, can still form an analogous viral IF4F (vIF4F) complex together with vIF4A and vIF4E. As recombinant expression of vIF4A and vIF4E from APMV yielded preparations unsuitable for biochemical characterization, we screened genomes of other giant DNA viruses of the *Mimiviradae* family for closely related proteins. We identified homologs of vIF4A, vIF4E, and vIF4G in cotonvirus^13^ (CJ, species *Cotonvirus japonicus*) that express to high levels and share ∼80% conserved amino acid identity with APMV and used these proteins in all subsequent biochemical assays. We found that vIF4G interacted with each protein individually and as a heterotrimeric complex, indicating these proteins assemble into an intact vIF4F complex (**Fig. 1G, Fig. S2B–C**). Notably, we further identified a short motif (YX_3_II) in vIF4G that shares cryptic homology with the canonical eIF4E binding motif (YX_4_LL) of human eIF4G but is repositioned proximal to the N-terminus (**Fig. S2E**)^14^. Mutating this motif to alanine led to reduced vIF4E binding, demonstrating that the mechanism of IF4F complex formation is broadly conserved (**Fig. 1H, Fig. S2D**). Together, we have identified highly divergent virus-encoded translation initiation factors that form a structurally conserved IF4F complex and interact with the host ribosome during infection.

### Viral IF4F is required for efficient replication

We next asked if the virally encoded IF4F complex is required for viral replication. We generated single gene knock-out viruses of each protein (*Δ4A, Δ4E*, and *Δ4G*) using homologous recombination, cloned them by limiting dilution, and genotyped each virus through diagnostic PCR (**Fig. 2A–B, Fig. S3A–B**). *Δ4A* and *Δ4G* viruses could only be rescued in amoebae stably expressing the respective viral translation factor, highlighting that these components are indispensable for production of infectious APMV particles. Loss of any individual component of the vIF4F complex severely diminished viral fitness, with ∼3–5-log reduction in viral titers compared to wild type (WT) APMV (**Fig. 2C–E**). Importantly, the effect on replication was specific to knockout of the IF4F proteins, as trans-complementation with the corresponding viral translation factor restored replication to WT levels. Notably, these results indicate the endogenous host eIF4F proteins cannot support viral replication in the vIF4F knockout strains, in agreement with substantial sequence divergence between host and viral translation factors that make it unlikely for key interaction surfaces to be compatible. Thus, giant virus-encoded translation factors are essential for viral replication and are functionally distinct from endogenous host eIF4F homologs.

**Figure 2.**
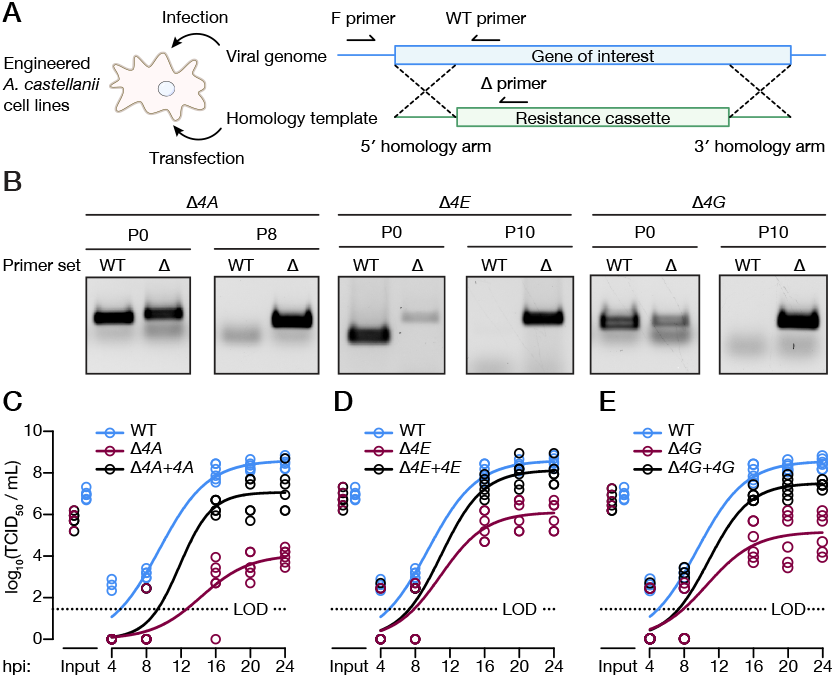
Viral IF4F is required for efficient replication. **A**. Schematic of constructs used for gene knockout (KO) by homologous recombination in engineered *A. castellanii*. The gene of interest (blue) is replaced by an antibiotic resistance cassette (green) through flanking homology arms. KO can then be verified by primers binding to the gene of interest (WT) or the resistance cassette (Δ). **B**. Diagnostic PCR of *Δ4A, Δ4E*, and *Δ4G* APMV after transfection (p0) and following passaging in the presence of selection agent (p8/p10). **C–E**. Viral growth curves of WT, *Δ4A, Δ4E*, and *Δ4G* APMV grown on WT or trans-complemented A. castellanii stable cell lines. Each circle represents one biological replicate (n = 6) with solid lines fitted by non-linear regression (logistic growth). The growth curve of WT APMV is replicated across all panels for visualization purposes. The limit of detection is indicated by a dotted line.

### Viral IF4F drives temporal and transcript-specific translation of structural proteins

The eIF4F complex plays a pivotal role in eukaryotic protein synthesis by regulating translation initiation. To define the molecular function of the vIF4F complex during viral replication we performed ribosome profiling and RNA-sequencing of amoebae infected with WT, *Δ4A, Δ4E*, or *Δ4G* APMV at early or late (4 or 8 hpi) timepoints. As expected, we observed a high degree of congruence between RNA-seq transcript abundances and ribosome-protected footprints (RPFs) across each sample and replicate (**Fig. S4A**). However, deletion of each subunit had no impact on total viral mRNA levels, indicating the vIF4F complex is not required for viral transcription or mRNA processing (**Fig. 3A**). In contrast, loss of vIF4F resulted in a severe decrease in late viral translation, with a ∼50% decrease in the ability of viral transcripts to be translated at 8 hpi but no effect at early points of replication (**Fig. 3B**). In agreement with a temporal regulatory role, viral IF4F mRNAs were only expressed at 8 hpi suggesting these proteins execute a dynamic switch to vIF4F-dependent translation initiation later in the infection cycle (**Fig. 3C**). Notably, when vIF4F was disrupted, metagene analysis of viral genes at 8 hpi showed reduced coverage by ribosomes at the start codon and an overall downward shift in the expression and translation of viral genes, consistent with a defect in translation initiation (**Fig. S4B–D**).

**Figure 3.**
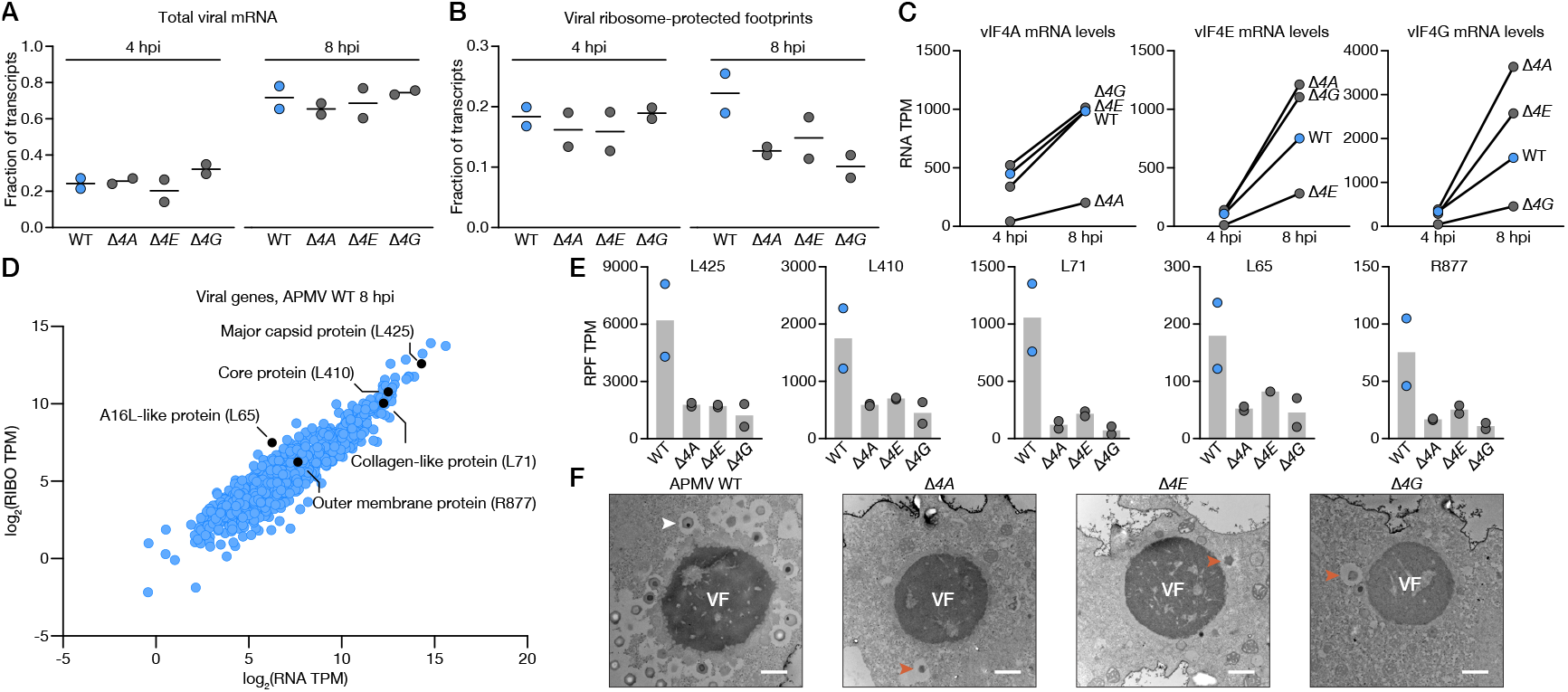
Viral IF4F drives temporal and transcript-specific translation of structural proteins. **A**. Normalized abundances of viral mRNA at 4 and 8 hpi from *A. castellanii* cells infected with WT, *Δ4A, Δ4E*, and *Δ4G* APMV. Each line represents the mean of two biological replicates. **B**. Normalized abundances of viral ribosome-protected footprints (RPFs) at 4 and 8 hpi from *A. castellanii* cells infected with WT, *Δ4A, Δ4E*, and *Δ4G* APMV. Each line represents the mean of two biological replicates. **C**. Normalized mRNA abundances of vIF4F subunits at 4 and 8 hpi. Each circle represents the mean of two biological replicates. In mutant viruses, reads assigned to the deleted gene largely map to the intact 5′ and 3′ UTRs. **D**. Normalized and log_2_-transformed RNA and RPF abundances of all viral transcripts detected at 8 hpi during APMV WT infection. A subset of key structural proteins are highlighted in black. Each dot represents the mean of two biological replicates. **E**. Abundance of RPFs corresponding to select viral structural transcripts. Each bar represents the mean of two biological replicates **F**. Transmission electron micrographs of *A. castellanii* cells infected with WT, *Δ4A, Δ4E*, and *Δ4G* APMV collected at 12 hpi. An example of an intact viral particle is indicated by a white arrow. Structurally malformed particles are indicated by orange arrows.

As replication proceeds, successful viral transmission relies on the ability to efficiently express proteins required for progeny virion formation and initiation of the subsequent infectious cycle. Correspondingly, we identified key late-expressed structural proteins^15^, such as the major capsid protein (L425), core protein (L410), collagen-like protein (L71), A16L-like protein (L65), and the outer membrane protein (R877), among the most highly translated viral mRNAs at 8 hpi (**Fig. 3D**). Importantly, the number of ribosome-protected footprints corresponding to these mRNAs, was reduced by 2–14-fold without an intact vIF4F (**Fig. 3E**). Knock-out of any vIF4F subunit led to reductions in translation efficiency (TE) for nearly all of these mRNAs except L65. We further confirmed that structural mRNAs could not associate with highly translating polysomes in the absence of vIF4F proteins (**Fig. S4E–G**). In support of the critical importance of vIF4F-mediated translation of these mRNAs, we observed a profound impact on virus assembly, as reflected in striking differences in transmission electron micrographs of WT versus vIF4F knockout APMV-infected amoebae. While all viruses formed fully developed viral factories (VFs) few, if any, viral particles budded from these replication sites in the absence of vIF4F components (**Fig.3F**). In sharp contrast, pseudo-icosahedral nascent virus particles were readily visible in WT APMV-infected cells in agreement with robust virus production (**Fig. 2C–E**). Therefore, giant virus translation factors provide dynamic regulation of protein synthesis and virion assembly, with a temporal switch to vIF4F-dependent initiation later in the infection cycle.

### The viral IF4F cap-binding subunit recognizes viral cap structures

The 5′ 7-methylguanosine cap (m^7^G) is a defining feature of eukaryotic mRNAs. As eIF4E recognizes mRNAs through coordination of this cap structure, m^7^G is critical for the first step of cap-dependent initiation of global protein synthesis in eukaryotes^16^. Both host and giant virus mRNAs exhibit this modification, leaving a canonical cap-binding protein with no basis to discriminate between the two. However, since the vIF4F complex is required for efficient translation of viral mRNAs, we hypothesized that features beyond m^7^G allow for specific recognition of viral mRNAs. The 5′ untranslated regions (UTR) of viral mRNAs exhibit strong sequence conservation not observed in host mRNAs, with a near invariable adenosine in the +1 positions of APMV mRNAs followed by AU-rich sequences (**Fig. 4A**). Given the conserved nucleotide identity proximal to the cap, we asked if viral IF4E evolved to exploit this difference and drive specific recognition of viral mRNA. We used differential scanning fluorimetry as an in vitro binding assay to analyze vIF4E binding to a diverse set of nucleotides that mimic the structure of mRNA 5′ caps. As expected, human eIF4E bound similarly to all ligands containing a 5′ m^7^G but did not interact with GTP. In contrast, vIF4E selectively interacted with a viral 5′ mRNA cap-mimic containing m^7^G followed by a 2′-O-methylated adenosine and bound poorly to m^7^GTP or any ligand with an unmethylated +1 nucleotide (**Fig. 4B**). Notably, giant viruses encode cap methyltransferases with homology to the eukaryotic capspecific mRNA (nucleoside-2′-O-)-methyltransferases (2′-OMTase) CMTR1 and CMTR2, which install 2′-O-methylation on the +1 and +2 nucleotides of m^7^G-capped transcripts^17^. In APMV, this 2′-OMTase (R383) is encoded immediately downstream of the trifunctional capping enzyme (R382), and therefore could coordinate the installation of 2′-O-methylation on the conserved +1 adenosine on viral mRNAs^18^. Indeed, such “cap-1” methylation of +1 adenosines is performed in megaviruses and poxviruses by a distinct class of viral 2′-OMTase^19^.

**Figure 4.**
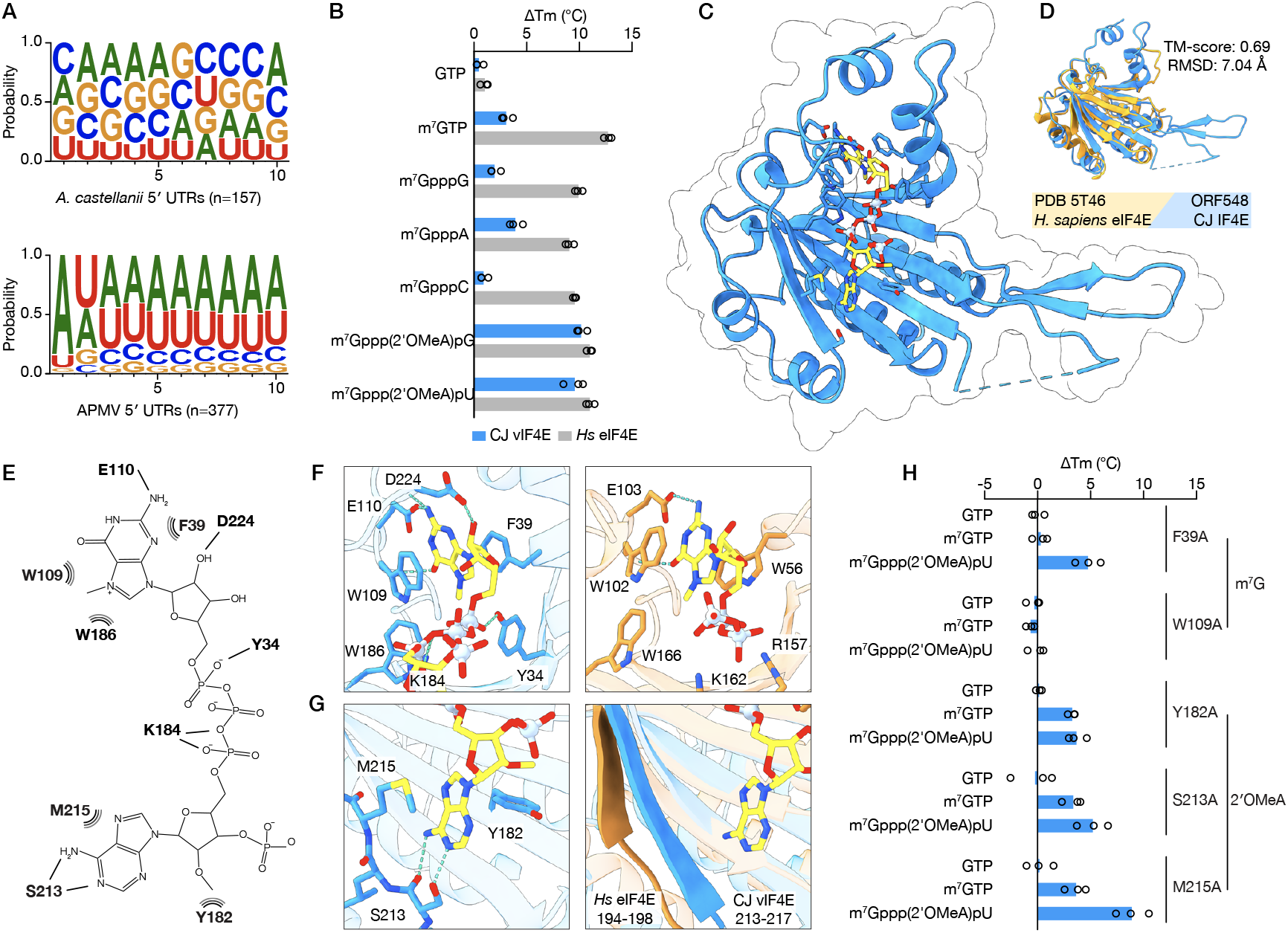
The viral IF4F cap-binding subunit recognizes viral cap structures. **A**. Weblogos of the first 10 nt of host (top) and viral (bottom) 5′ UTRs. **B**. ΔTm values of *Hs*. eIF4E (grey bars) and cotonvirus (CJ) vIF4E (blue bars) upon binding to nucleotide mimics of 5′ cap structures. Each bar represents the mean of three biological replicates. **C**. X-ray crystal structure of CJ vIF4E in complex with m^7^Gppp(2′OMeA)pU at 2.26 Å (PDB 9E5B). **D**. Structural comparison of CJ vIF4E and *Hs*. eIF4E (PDB 5T46) generated using Foldseek. **E**. Interaction diagram highlighting the residues that coordinate m^7^Gppp(2′OMeA)pU binding. Dashed teal lines indicate hydrogen bonds while curved black lines indicate π-stacking and VdW forces. **F**. Detailed view of the m^7^G binding site in CJ vIF4E (left) and *Hs*. eIF4E (right). **G**. Detailed view of the 2′OMeA binding site in CJ vIF4E (left) and the corresponding region in *Hs*. eIF4E (right). **H**. ΔTm values of CJ vIF4E with mutations to putative m^7^G or 2′OMeA coordinating residues. Each bar represents the mean of three biological replicates. All residue numbers correspond to CJ vIF4E.

2′-O-methylation at the +1 position of mRNAs is known to increase transcript stability and translation, but is also a key aspect of the discrimination between self and nonself mRNAs in eukaryotes that allows for initiation of antiviral programs^20–22^. Therefore, to define the mechanism of specific viral mRNA recognition by vIF4F, we next determined a 2.3 Å crystal structure of vIF4E in complex with m^7^Gppp(2′OMeA)pU (**Fig. 4C**). The structure of vIF4E reveals core **features shared with huma**n eIF4E in addition to novel adaptations that enable viral mRNA selection through unique interactions with the ligand (**Fig. 4D–E, Fig. S5D**)^16,23^. Human eIF4E, and other canonical type 1 eIF4E proteins, binds m^7^G in a concave binding pocket that coordinates hydrogen bonds with a conserved glutamic acid and pi-stacking with two conserved tryptophan residues forming an aromatic sandwich. An additional tryptophan reads out the methylation status of the guanosine (**Fig. 4F, right**). In vIF4E, the m^7^G ligand is recognized in a similar binding pocket through hydrogen bonds with the well-conserved E110, but the aromatic sandwich instead consists of W109 and a conserved F39 contributing to a slightly tilted geometry of the binding pocket relative to human eIF4E (**Fig. 4F, left**). These differences in the m^7^G binding pocket explain why vIF4E is a relatively weak m^7^G-binder and are reminiscent of the atypical binding pockets of type 2 and 3 eIF4E proteins, which are proposed to regulate eukaryotic translation on a transcript-specific rather than global level and have been implicated in adaptation to stress^24,25^. At the base of vIF4E, 2′OMeA is coordinated by three conserved residues–S213, M215, and Y182. Changing the identity of the +1 nucleobase would disrupt the hydrogen bonds to S213, explaining the reduced affinity of vIF4E to cap mimics lacking a +1 A (**Fig. 4G, left**). The corresponding region in human eIF4E is not positioned to make such contacts with a nucleotide, and the entire ligand-binding groove of human eIF4E is more open than that of vIF4E (**Fig. 4G, right, Fig. S5A–B**). We biochemically tested cap-binding of a panel of vIF4E mutant proteins using differential scanning fluorimetry and confirmed that vIF4E has evolved novel adaptations to recognize both m^7^G and 2′OMeA. Mutations that disrupt the m^7^G binding site (F39A, W109A) were detrimental to cap binding, but the F39A mutant retained residual binding specificity to a cap mimic containing 2′OMeA. Conversely, when the site of 2′OMeA recognition was mutated (Y182A, S213A) specific recognition of m^7^Gppp(2′OMeA)pU was abolished while binding to m^7^G was unaffected (**Fig. 4H**). M215A did little to impact vIF4E cap-binding, possibly due to the conservative nature of the M-to-A substitution. To validate these biochemical findings, we generated *A. castellanii* cell lines expressing APMV vIF4E with the corresponding mutations to the m^7^G binding site (W108A), 2′OMeA binding site (S230A), or vIF4G with a disrupted vIF4E:vIF4G interface (vIF4G-ALA). Unlike the WT counterparts, these mutant proteins failed to support replication of APMV *Δ4E* and *Δ4G*, demonstrating the functional relevance of these residues and underscoring the conserved nature of the ligand binding interfaces identified in the purified CJ vIF4E homolog (**Fig. S5C**). vIF4E thus couples canonical m^7^G cap-binding properties with a novel mode of mRNA sequence and modification-dependent recognition, indicating that vIF4E has evolved to integrate regulatory functions distributed across several proteins in eukaryotes. Altogether, our results illustrate how evolutionary innovations to the IF4F complex are sufficient to drive the emergence of transcript-specific translation regulation, suggesting similar additive adaptations could have contributed to the complex specialized translation observed in eukaryotes.

### Expression of viral IF4F by mimivirus confers resistance to abiotic cell stresses

The long-standing dogma is that viruses share a universal dependence on host cell translation factors and machinery for viral protein synthesis. What evolutionary forces drive giant DNA viruses to encode a eukaryotic-like translation initiation factor complex? Notably, these viruses infect diverse unicellular protists, including amoeba, choanoflagellates, and ciliates, which experience unique ecological pressures that differ from those of mammalian hosts of well-characterized viruses. For example, protists frequently encounter dynamic and stressful environments, including fluctuating pH or temperature levels and desiccation. Indeed, giant DNA virus sequences have been isolated from extreme environments like the Siberian permafrost and geothermal springs^26,27^. Thus, these viruses must be able to replicate despite large physiological shifts in the host cellular state. In eukaryotes, exposure to diverse abiotic stresses triggers general translation shutoff while simultaneously activating specialized translation initiation pathways that promote the synthesis of proteins to help return cellular homeostasis^28–32^. We therefore asked if vIF4F might function in a manner analogous to specialized translation pathways to provide efficient translation initiation under conditions when host translation is impaired. We subjected amoebae to a set of conditions that mimic relevant environmental stresses encountered in their complex ecosystem and measured the effect on APMV replication. Strikingly, while WT APMV replication was highly resistant to nutrient deprivation, thapsigargin-induced endoplasmic-reticulum (ER) stress, or menadione-induced oxidative stress, viruses lacking an intact vIF4F exhibited additional 1–2 log defects in virus production compared to baseline host conditions (**Fig. 5A, Fig. S5F**). Cold shock at 18°C had a uniformly negative effect on viral replication, indicating that certain stress pathways cannot be circumvented by vIF4F activity. As a comparison, we utilized noumeavirus (NMV), a related virus with multiple features similar to APMV. Both viruses belong to the same taxonomic class^33^, infect the same host^34^, and exhibit similar genomic and structural features, including AU-rich intergenic regions with conserved motifs^35,36^, icosahedral virion morphology^34^, and an exclusively cytoplasmic replication cycle organized around a viral factory formed via liquid-liquid phase separation^37^. Notably, NMV does not encode any vIF4F subunits, yet it shares a similar signature of late-expressed proteins with APMV, including the capsid protein and 24% of the 50 most highly expressed late genes (**Table S2**). NMV did not exhibit the same replication plasticity, suggesting that although vIF4F is not universally required for giant DNA virus replication, its presence in APMV may enable viral advantages in conditions of cellular stress (**Fig. 5A, Fig. S5E**). To directly visualize how APMV alters translation during stress we generated polysome profiles from infected and mock-infected cells under starvation or ER stress. While mock-infected cells show a loss of polysomes and concordant increase in monosomes as expected with stress-induced shutoff of general protein synthesis and a switch to specialized translation, profiles from APMV-infected cells remain largely unaltered under these restrictive conditions (**Fig. 5B–C**). Intriguingly, while polysome fractionation and qRT-PCR verified that key viral transcripts remain associated with polysomes during starvation, this effect is muted during ER stress, suggesting that viral translation during stress is not uniform. Additionally, unlike most viral infections that typically trigger global translation shutoff, APMV infection results in minimal redistribution of translation complexes from polysomes to monosomes (**Fig. 1B, Fig. S1D**). Altogether, these findings suggest that translation during giant virus infection has been reprogrammed to broadly bypass host machinery shutoff mechanisms typically conserved across infection or stress^38,39^. Collectively, our results suggest that the evolution of viral translation factors contributes to enhanced fitness during host stress and allows these viruses to replicate under conditions that are detrimental to most mammalian viruses.

**Figure 5.**
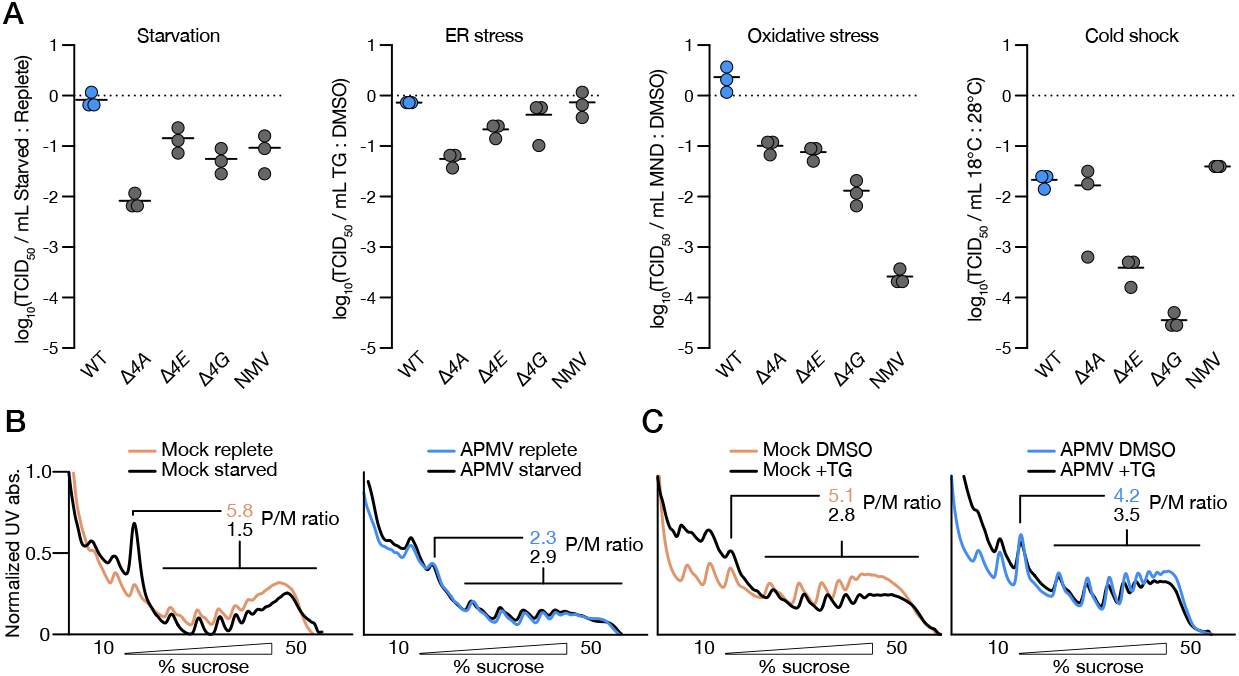
Expression of viral IF4F by mimivirus confers resistance to abiotic cell stresses. **A**. Relative viral titers of WT, *Δ4A, Δ4E, Δ4G* APMV and NMV under starvation, ER-stress, oxidative stress, and cold shock. Each line represents the mean of three biological replicates. **B**. Representative polysome profiles from APMV- and mock-infected cells under replete and starvation conditions. **C**. Representative polysome profiles from APMV- and mock-infected cells under TG or DMSO treatment.

## Discussion

The evolution of complex life is closely linked to the emergence of mechanisms that regulate protein synthesis to finely tune the composition of the proteome. Key among these mechanisms has been the sequential expansion of translation factors, which regulate ribosome function to control protein synthesis in a temporal, localization, and RNA-specific manner^40–42^. Here, through the discovery and characterization of a eukaryotic translation initiation complex in viruses, we expand this paradigm of cellular life to also include viruses. The viral IF4F cap-binding complex coordinates a temporal switch to enable translation of structural proteins late in infection, and this transcript-specific regulation occurs through evolutionary innovations to m^7^G cap recognition. We further show that the viral IF4F complex allows for resilient replication under stress conditions, demonstrating that dynamic regulation of protein synthesis mediated by translation factors surprisingly is a feature shared by cellular organisms and viruses. By uncovering a fundamentally distinct form of viral translation, these findings establish giant DNA viruses as a model to discover new modes of translation regulation and further define the boundaries between cellular life and viruses.

In eukaryotes, eIF4F acts to regulate initiation of global protein synthesis based on recognition of the m^7^G cap. In contrast, viral IF4F has adapted several evolutionary novelties that promote specialized viral gene expression and fitness. First, vIF4E has gained selectivity for viral mRNAs by making additional contacts with the near invariable 2′-O-methylated adenosine at the +1 position. Given this shift in selectivity to elements beyond the m^7^G cap, it will be of interest to understand which mode of cap recognition has a more ancient role in translation regulation by further investigating the evolutionary origins of viral IF4F. Intriguingly, the crystal structure of vIF4E also reveals two loops (β3–β4 and β7–β8) that extend from the cap-binding pocket in a position that could make additional downstream contacts with the AU-rich viral 5′ UTRs and provide additional ligand specificity or regulatory functions. Second, while viral and eukaryotic IF4A display high sequence conservation in the DEAD-box helicase motifs, vIF4A contains virus-specific N-terminal and internal extensions of unknown function (Fig. S1A-C). In the canonical model of translation initiation, eIF4A unwinds secondary structure of long and complex 5′ UTRs to facilitate efficient scanning by the 48S initiation complex. However, as the viral 5′ UTRs are short and unstructured, it is puzzling why vIF4A is required for viral replication and protein synthesis. Recent findings suggest eIF4A acts to remove mRNAs bound to IF4F at unproductive cap-distal sites^43^. It will therefore be of interest in future studies to investigate if vIF4A acts in accordance with this model, or if the virus-specific adaptations in vIF4A, similarly to vIF4E, tune the ability of viral IF4F to be selective for viral mRNAs. Third, despite lacking several domains essential for orchestrating eukaryotic translation initiation, vIF4G successfully coordinates vIF4F complex formation and ribosome association. Determining how a minimal IF4F complex is recruited to host ribosomes will have significant implications for understanding the origin and evolution of eukaryotic translation initiation. Altogether, towards solving the long-standing enigma of why giant viruses encode large complements of cellular genes that are absent from the rest of the virome, our findings suggest these are sufficient to enable complex viral gene expression regulation that is traditionally a hallmark of cellular life.

Our finding of parallel IF4F translation pathways in viruses and eukaryotes also raises the question of whether viruses contributed to the expansion of transcript-specific translation mechanisms important in specializing the function of eukaryotic cells. While viruses are pathogens, they also drive molecular evolution across all domains of life. We speculate that the ability of viral infection to promote protein synthesis plasticity in response to environmental and intrinsic stresses could act as a factor to shape the evolution of cellular plasticity. Indeed, giant DNA viruses also harbor genes related to photosynthesis, DNA maintenance/repair, energy metabolism, and several biosynthetic pathways. Thus, viral evolution could more broadly shape biological and ecological community dynamics.

### Limitations of the study

Here, we combine in vitro, bulk-cell translation, and viral replication assays to discover and define the functions of vIF4F. However, increasing evidence indicates that translation during APMV infection is spatially organized with viral mRNAs, tRNAs, and translation factors colocalizing in the zone immediately surrounding the VF^37,44,45^. This segregation of translation components could contribute to viral translation regulation and selectivity which we do not capture using assays with whole-cell lysates described herein. Indeed, the localized nature demonstrated for some viral translation factors also raises the question if there are specialized circumstances when the viral and host translation factors interact. Since the highly divergent viral IF4F subunits were exclusively detected without an intact host eIF4F in translation complexes from infected cells, we believe the formation of chimeric IF4F complexes is unlikely under normal infection conditions. However, there could be other ecological or biotic conditions where such interactions are critical or occur in the cell in a localized manner. Future studies leveraging sub-cellular fractionation of infected cells and identification of VF-associated translation complexes are of interest to bridge this gap. Finally, we note that work on giant DNA viruses and non-model organisms such as *Acanthamoeba spp*. is limited by the lack of well-annotated genomes and transcriptomes. For example, the relatively small number of annotated host 5′ UTRs limits our current ability to determine how the identity of the +1 nucleotide impacts translation of host mRNAs, and the complete lack of annotated NMV 5′ UTRs precludes direct comparisons to APMV. In the future, improved annotations via comparative genomics or experimental validation of transcription start sites will be of great value to studying evolution of protein synthesis in these remarkable viruses.

## Acknowledgements

The authors thank the Taplin Mass Spectrometry Facility for proteomics services; K. Chat and members of the Lee and Kranzusch labs for discussions. Electron Microscopy imaging, consultation, and services were performed in the HMS Electron Microscopy Facility. This work was funded by grants to A.S.Y.L. from the Pew Biomedical Scholars Program, The G. Harold and Leila Y. Mathers Charitable Foundation, and the National Institutes of Health (1R35GM142527); grants to P.J.K. from The G. Harold and Leila Y. Mathers Charitable Foundation, the Pew Biomedical Scholars program, the Parker Institute for Cancer Immunotherapy and the National Institutes of Health (no. 1DP2GM146250); grants to C.A. from the European Research Council (ERC) under the European Union’s Horizon 2020 research and innovation program (832601); an NRSA fellowship to R.H. (F31NS132412). H.B is funded by the European Union (ERC, ViDaMa, grant agreement No. 101160452, ERC-2024-STG). Views and opinions expressed are however those of the author(s) only and do not necessarily reflect those of the European Union or the European Research Council. Neither the European Union nor the granting authority can be held responsible for them. J.M.F. is supported by the Branco Weiss Fellowship and the Moderna Global Fellowship.

## Author contributions

Conceptualization, JMF, ASYL, PJK; Investigation, JMF, ABH, RH, JMG; Resources, HB, CA; Data Curation, JMF, ASYL; Writing – Original Draft, JMF, ASYL; Writing – Review & Editing, JMF, ABH, RH, HB, CA, PJK, ASYL; Visualization, JMF, ABH; Supervision, ASYL, PJK; Funding Acquisition, JMF, ASYL, PJK.

## Materials and Methods

### Resource Availability

Further information and requests for resources and reagents should be directed to and will be fulfilled by the lead contact, Amy S.Y. Lee.

### Materials Availability

Reagents and materials produced in this study are available from the lead contact pending a completed Materials Transfer Agreement.

### Data and Code Availability

- All sequencing data have been deposited in the Gene Expression Omnibus under accession number GSE290007 and are publicly available as of the date of publication.
- Coordinates of the vIF4G and vIF4E in complex with m^7^Gppp(2′OMeA)pU structures have been deposited to the Protein Data Bank (PDB) under the accession number 9E5A and 9E5B.
- This paper does not report original code.
- Any additional information required to reanalyze the data reported in this paper is available from the lead contact upon request.

### Cells and viruses

*Acanthamoeba castellanii* str. Neff (ATCC, 30010) were maintained at 28°C in PYG (20 g L^−1^ peptone, 2 g L^−1^ yeast extract, 9 g L^−1^ glucose, 4 mM MgSO_4_, 0.4 mM CaCl, 50 µM Fe(NH_4_)_2_(SO_4_)_2_, 2.5 mM Na_2_HPO_4_, 2.5 mM KH_2_PO_4_, 4 mM sodium citrate) supplemented with 1% (v/v) penicillin-streptomycin (Gibco, 15140122). Cells were sub-cultured at 2–3-day intervals to maintain the cells in trophozoite stage. The mimivirus isolate used herein (APMV, species *Mimivirus bradfordmasseliense*) was derived from the first mimivirus isolate^9^. The noumeavirus isolate used herein (NMV, species unassigned) was derived from the first noumeavirus isolate^34^ Purified mimivirus and noumeavirus preparations were prepared as previously described^46^. Briefly, ∼20 million Acanthamoeba castellanii cells cultured in PYG were inoculated at a multiplicity of infection (MOI) of 0.01. Cells were monitored for cytopathic effect (CPE) which peaked at 72 hours postinfection (hpi). Supernatants were then harvested and clarified by low-speed centrifugation to remove cellular debris. Viral supernatants were overlaid onto a 10 mL cushion of 24% (w/v) sucrose solution and purified by ultracentrifugation at 36,000 × g in an SW-32 Ti rotor for 30 min at 4°C. Pellets were resuspended on ice in PBS before aliquoting and storing at −80°C. All viral titers were determined by the TCID_50_ method. *A. castellanii* grown in PYG were seeded in 96-well plates at a density of 1,000 cells per well. For titering, viral supernatants were serially diluted in PYG and then added to cells. Cells were monitored for CPE and scored as positive or negative at 72 hpi. TCID_50_ and a 95% confidence interval were calculated using the improved Kärber method as previously described^47^. TCID_50_ values less than the limit of detection were set to 0. To generate viral growth curves, WT *A. castellanii* or *A. castellanii* trans-complemented with vIF4A, vIF4E or vIF4G were grown in PYG to a confluency of ∼90%. Cells were then inoculated with WT, *Δ4A, Δ4E*, and *Δ4G* APMV at an MOI of 5. The inoculum was adsorbed for 1 h before aspirating and washing the cells three times with PYG. Supernatants were collected at the indicated times postinfection and titered by TCID_50_.

### Generation of trans-complementing cell lines

Trans-complementing cell lines were generated as previously described^48^. Briefly, *A. castellanii* grown in complete PYG were transfected with plasmids encoding vIF4A, vIF4E, vIF4G, vIF4E-W108A, vIF4E-S230A or vIF4G-ALA, followed by a neomycin or nourseothricin resistance cassette. 10 µg of DNA and 15 µg of PEI 40K (Polysciences, 24765) were separately diluted in 100 µl of plain PYG. DNA and PEI dilutions were then combined and incubated at room temperature for 20 min. The volume of the transfection reaction was adjusted to 500 µl before dropwise adding the mixture to the cells. Transfection was allowed to proceed for 14 h before washing the cells once with plain PYG. At 24 h post-transfection the media was replaced with complete PYG containing 30 µg mL^−1^ of Geneticin/G-418 (Gibco, 10131035). Cells were passaged under selective pressure until a fully resistant population remained.

### Generation of APMV knockout viruses

Knock-out viruses were generated as previously described^48^. Briefly, to generate APMV *Δ4E* and *Δ4G, A. castellanii* trans-complemented with vIF4E or vIF4G were infected with WT APMV at an MOI of 5. At 1 hpi cells were washed with PBS five times before transfection with PCR amplicons containing 5′and 3′ homology arms flanking a neomycin or nourseothricin resistance cassette. 10 µg of DNA and 15 µg of PEI 40K (Polysciences, 24765) were separately diluted in 100 µl of PYG. DNA and PEI dilutions were then combined and incubated at room temperature for 20 min. The volume of the transfection reaction was adjusted to 500 µl before dropwise adding the mixture to the cells. At approximately 16 hpi CPE was readily observed and supernatants were harvested. To generate APMV *Δ4A, A. castellanii* transcomplemented with vIF4A were transfected as above. Transfected cells were incubated for approximately 6 h and were then infected with WT APMV at an MOI of 5. At 1 hpi cells were washed with PBS five times and then incubated until CPE was readily observed at which point supernatants were harvested. Successful homologous recombination was verified using diagnostic PCR with mutant- or WT-specific primer pairs. 5 µl of virus-containing supernatants was added directly to Q5 polymerase (NEB, M0491S) reactions to generate ∼300 nt long amplicons which were visualized on 1.5% agarose gels. Supernatants containing recombinant viruses were then passaged on trans-complemented *A. castellanii* cells in the presence of 100–150 µg mL^−1^ nourseothricin (*Δ4A* and *Δ4G*) or 25–50 µg mL^−1^ G-418/geneticin (*Δ4E*). After multiple passages followed by diagnostic PCR, limiting dilution was used to generate fully recombinant viral populations. Viral supernatants were serially diluted in PYG before infecting trans-complemented *A. castellanii* cells in a 96-well format. Wells at the highest dilution with clear signs of CPE at 72 hpi were screened by diagnostic PCR and pure APMV *Δ4A, Δ4E*, and *Δ4G* stocks were then generated as described above by growing mutant viruses on transcomplemented *A. castelanii* in the absence of any selection agent.

### Polysome profiling

*A. castellanii* grown to ∼90% confluency in 150 cm^2^ flasks were mock-infected or infected with WT APMV at a MOI of 3. At 1 hpi the inoculum was aspirated and cells were washed once with PYG. At 8 hpi, 10 mL of icecold amoeba salt solution (4 mM MgSO_4_, 0.4 mM CaCl, 50 µM Fe(NH_4_)_2_(SO_4_)_2_, 2.5 mM Na_2_HPO_4_, 2.5 mM KH_2_PO_4_, 4 mM sodium citrate) supplemented with 9 g L^−1^ of glucose and 100 µg mL^−1^ of cycloheximide were added to each flask. Cells were then harvested by scraping, pelleted by low-speed centrifugation, washed once in amoeba salt solution, and flashfrozen in liquid nitrogen. Cell pellets were lysed in lysis buffer (20 mM Tris-HCl pH 7.5, 200 mM NaCl, 10 mM MgCl_2_, 0.1% (v/v) Triton X-100, 1 mM DTT, 100 µg mL^−1^ cycloheximide) and triturated through a 21-gauge needle five times. Lysates were clarified by centrifugation at 10,000 × g for 5 min at 4°C. The RNA concentration of each lysate was measured and 1,000 A_260_ units of lysates were layered on 10–50% (w/v) sucrose gradients in polysome buffer (20 mM Tris-HCl pH 7.5, 150 mM NaCl, 10 mM MgCl_2_, 0.2 mM spermidine, 1 mM DTT). Gradients were centrifuged at 35,000 rpm for 2 h at 4°C in a Beckman SW 41 Ti rotor and separated using a Brandel gradient fractionator with UV absorbance at 260 nm monitored by a Dataq system. UV traces were manually trimmed and then imported to QuAPPro^49^. Baseline adjustment, normalization, and a 0.2 smoothing factor were applied before alignment of each trace using the 80S peak as the anchor point.

### Mass spectrometry of translation complexes

Fractions corresponding to 40S subunits were collected from polysome profiles prepared as described above. The fractions were concentrated using 30 kDa MWCO centrifugal filters to ∼30 µL and then proteins were run ∼3 mm into a 2–20% gradient gel (Bio-Rad, 4561094) before staining with Coomassie stain. Bands were excised and cut into approximately 1 mm^3^ pieces. Gel pieces were then subjected to a modified in-gel trypsin digestion procedure^50^. Gel pieces were washed and dehydrated with acetonitrile for 10 min followed by removal of acetonitrile. Pieces were then completely dried in a speed-vac. Rehydration of the gel pieces was performed with 50 mM ammonium bicarbonate solution containing 12.5 ng µL^−1^ modified sequencing-grade trypsin (Promega, Madison, WI) at 4°C. After 45 min the excess trypsin solution was removed, replaced with 50 mM ammonium bicarbonate solution to cover the gel pieces, and samples were then placed at 37°C overnight. Peptides were later extracted by removing the ammonium bicarbonate solution, followed by one wash with a solution containing 50% acetonitrile and 1% formic acid. The extracts were then dried in a speed-vac for ∼1 h and samples were stored at 4ºC until analysis. On the day of analysis, the samples were reconstituted in 5–10 µL of HPLC solvent A (2.5% acetonitrile, 0.1% formic acid). A nano-scale reverse-phase HPLC capillary column was created by packing 2.6 µm C18 spherical silica beads into a fused silica capillary (100 µm inner diameter × ∼30 cm length) with a flame-drawn tip^51^. After equilibrating the column each sample was loaded via a Famos autosampler (LC Packings, San Francisco CA) onto the column. A gradient was formed and peptides were eluted with increasing concentrations of solvent B (97.5% acetonitrile, 0.1% formic acid). As peptides eluted, they were subjected to electrospray ionization and then entered a Velos Orbitrap Pro ion-trap mass spectrometer (Thermo Fisher Scientific, Waltham, MA). Peptides were detected, isolated, and fragmented to produce a tandem mass spectrum of specific fragment ions for each peptide. Peptide sequences (and hence protein identity) were determined by matching protein databases with the acquired fragmentation pattern by the software program Sequest (Thermo Fisher Scientific, Waltham, MA)^52^. All databases include a reversed version of all the sequences and the data was filtered to between a one and two percent peptide false discovery rate. The relative abundance of each detected protein was normalized in two steps. First, the sum intensity of detected peptides mapped to each protein was normalized by its molecular weight. Second, the normalized peptide intensity for a given protein was divided by the total sum intensity of each sample and log_2_-transformed to allow for cross-sample comparisons of abundances.

### Structural prediction and homology searches

The Colabfold implementation of AlphaFold2 was used to derive structural models of orphan viral proteins identified in 40S fractions from infected cells^53^. For each protein the model with the highest pLDDT score was selected as a Foldseek query to search for structural homologs^54^. The resulting list of hits was then manually searched for translation-related genes. For the structural comparison between vIF4G and eIF4G the top hit from a search against the PDB100 database (PDB 8J7R) was selected. For the structural comparison between vIF4E and eIF4E the third hit from a search against the PDB100 database (PDB 5T46) was selected since it is the first human protein and shares virtually all statistics with the first two hits.

### Phylogenetic analysis and protein alignments

APMV vIF4G (UniProt ID Q5UPT9) and Klosneuvirus vIF4G (UniProt ID A0A1V0SJ99) were used as queries for separate blast-p searches for viral homologs, which after filtering out spurious hits resulted in 23 homologs. An AlphaFold2 model of APMV vIF4G was also used as a Foldseek query against the AFDB-Proteome database, which resulted in 52 hits after filtering for >99% probability. The combined set of viral and eukaryotic vIF4G homologs were aligned using Clustal Omega implemented in Geneious Prime^55^ and phylogenetic trees with branch supports were generated using FastTree^56,57^. Sequence alignments of vIF4A, vIF4E, and vIF4G were generated using Clustal Omega implemented in Jalview with default settings^58^. Alignments were colored by BLOSUM62 with a conservation threshold score set at 30.

### Gene ontology enrichment analysis

Host proteins identified in MS/MS data from mock-infected and APMV WT-infected conditions were used as inputs for GO enrichment analysis using ShinyGO^59^. FDR cutoff was set to 0.05 with a pathway minimum size of 2. The 20 most highly enriched pathways under each condition are displayed and colored by FDR.

### Protein expression and purification

Constructs were cloned by Gibson assembly into a custom pET vector allowing for expression of a 6×His-SUMO2 fusion protein^60^. Plasmids were transformed into either the E. coli strain BL21-DE3 RIL (Agilent, 230245) or LOBSTR-BL21(DE3)-RIL (Kerafast, EC1002). Transformed E. coli were plated onto 1.5% agar plates containing MDG media (0.5% (w/v) glucose, 2 mM MgSO_4_, 25 mM Na_2_HPO_4_, 25 mM KH_2_PO_4_, 50 mM NH_4_Cl, 5 mM Na_2_SO_4_, 0.25% (w/v) aspartic acid, 2–50 µM trace metals, 100 µg mL^−1^ ampicillin and 34 µg mL^−1^ chloramphenicol) and grown at 37°C overnight. Three colonies were then inoculated into 30 mL of MDG media and were grown overnight at 37°C with shaking at 230 rpm. 10 mL of the overnight culture was added to 1 L of M9ZB media (47.8 mM Na_2_HPO_4_, 22 mM KH_2_PO_4_, 18.7 mM NH_4_Cl, 85.6 mM NaCl_2_, 1% (w/v) Casamino Acids (ThermoFisher, 223110), 0.5% (v/v) glycerol, 2 mM MgCl_2_, 2–50 µM trace metals, 100 µg mL^−1^ ampicillin, and 34 µg mL^−1^ chloramphenicol) and were grown at 37°C with shaking at 230 rpm until the optical density at 600 nm reached above 1.5. Cultures were then cooled on ice, 0.5 mM IPTG was added, and cultures were grown at 16°C with shaking at 230 rpm. Cells were pelleted by centrifugation at 2,400 × g for 15 min, resuspended in lysis buffer (20 mM HEPES-KOH pH 7.5, 400 mM NaCl, 10% (v/v) glycerol, 30 mM imidazole pH 7.0, 1 mM DTT), and lysed by sonication. Lysates were cleared by centrifugation at 20,000 rpm for 30 min and the soluble fraction was mixed with 8 mL equilibrated Ni-NTA agarose (Qiagen, 30250). The solution was transferred to a gravity filtration column and the flow through was re-applied to the column. The resin was washed with 20 mL of lysis buffer, 70 mL of wash buffer (20 mM HEPES-KOH pH 7.5, 1 mM NaCl, 10% (v/v) glycerol, 30 mM imidazole, 1 mM DTT), and 35 mL of lysis buffer. Proteins were eluted with 20 mL of elution buffer (20 mM HEPES-KOH pH 7.5, 400 mM NaCl, 10% (v/v) glycerol, 300 mM imidazole, 1 mM DTT). Eluted proteins were mixed with human SENP2 protease and then dialyzed in 1.75 L of dialysis buffer overnight (20 mM HEPES-KOH pH 7.5, 250 mM KCl, 1 mM DTT). Dialyzed protein samples were concentrated to ∼1.5 mL with 30 kDa MWCO centrifugal filters. Samples were then applied to a 16/600 Superdex 75 column equilibrated with a storage buffer (20 mM HEPES-KOH pH 7.5, 250 mM KCl,1 mM TCEP). Fractions containing protein were collected, pooled, and concentrated to >10 mg mL−1 before flash freezing in liquid nitrogen for storage.

### Crystallography

For APMV vIF4G (54-312AA), samples at 36 mg mL^−1^ in storage buffer were diluted to 5 mg mL^−1^ in 20 mM HEPES-KOH pH 7.5 and 1 mM TCEP. Initial screening was performed in a hanging drop format with 96-well trays with 70 µl reservoir by mixing 200 nL of protein solution with 200 nL of well solution with a Mosquito instrument (SPT Labtech). Further optimization was performed with 15-well trays with a 350 µL reservoir solution by mixing 1 µL of protein solution with 1 µL of reservoir solution. Crystals formed with a reservoir solution of 0.2 M NaSCN and 20% PEG 3350 (w/v) after ∼2 days. The crystals were dipped into reservoir solution supplemented with 20% (v/v) ethylene glycol and harvested with a 0.2 µm nylon loop. X-ray diffraction data were collected at Diamond Light Source MX beamline. Data was processed using autoProc^61^. Experimental phase information was determined by molecular replacement using Phaser using a model predicted by AlphaFold2. Structural modeling was done using Coot^62^ and refinement was done in Phenix^63^. All structural figures were generated using ChimeraX^64^. For CJ vIF4E (1-236AA), samples at 56 mg mL^−1^ in storage buffer were diluted to 10 mg mL^−1^ in 20 mM HEPES-KOH pH 7.5, 1 mM TCEP, and 60 mM KCl and supplemented with 0.5 mM CleanCap Reagent AU (TriLink, N-7114). The Protein/ligand mixture was incubated for 25 min on ice before it was used in crystallization experiments. Screening and optimization were performed as described above. The best crystals were obtained in a solution of 2 M NaCl and 12% (w/v) PEG 6000 after ∼1 day. To harvest, the crystals were dipped into a solution containing 1.9 M NaCl, 12% (w/v) PEG 6000, 0.5 mM CleanCap Reagent AU, and 22% (v/v) ethylene glycol with a 0.2 µm nylon loop. X-ray diffraction data were collected at National Synchrotron Light Source II Blank beamline. Data was processed using autoProc^61^. Experimental phase information was determined by molecular replacement in Phaser using a model predicted by AlphaFold2. Structural modeling was done with Coot^62^ and refinement was done in Phenix^63^. All structural figures were generated with ChimeraX^64^. The structure of APMV vIF4G is available under the PDB accession number 9E5A. The structure of CJ vIF4E is available under the PDB accession number 9E5B.

### vIF4F pulldown experiments

For tagged proteins, constructs were cloned into a custom pET16 vector encoding a 6×His fusion protein using restriction enzyme cloning. All proteins were expressed as described above. vIF4A (YP_010841689.1), vIF4E (YP_010841645.1), and vIF4G (YP_010841617.1) from cotonvirus were purified, as described above, by Nickel-NTA purification. Proteins were then dialyzed, concentrated to >10mg mL^−1^, and stored in dialysis buffer (20 mM HEPES-KOH pH 7.5, 250 mM KCl, 1 mM DTT) with the exception of 6×His-vIF4E which was stored in dialysis buffer supplemented with 10% (v/v) glycerol. For pulldowns, proteins were combined and carefully diluted in binding buffer (20 mM HEPES-KOH pH 7.5, 50 mM KCl, 10 mM imidazole, 1 mM TCEP) to 200 µL to achieve a final salt concentration of 50 mM KCl and final protein concentrations of 5 µM of 6×His-tagged protein and 15 µM of untagged protein. Proteins were incubated on ice for 10 min to allow complex formation. 10 µL of each sample was saved to visualize the input and the rest was added to 50 µL of Ni-NTA agarose resin in a size exclusion mini spin column (Epoch, 2060-250) and incubated for 2 min. The solution was centrifuged to remove liquid and flow through was discarded. The resin was washed 3 times with 500 µL of binding buffer (20 mM HEPES-KOH pH 7.5, 50 mM KCl, 10 mM imidazole, 1 mM TCEP). Protein was eluted with elution buffer (20 mM HEPES, 400 mM KCl, 300 mM imidazole, 10% (v/v) glycerol, 1 mM TCEP) and 5 µL of each condition was visualized by SDS-PAGE using 15% acrylamide gels and Coomassie staining.

### Ribosome profiling and bioinformatics analyses

Lysates for ribosome profiling were generated from infected and mock-infected *A. castellanii*. All cells were cultured in 150 cm^2^ flasks in complete PYG until ∼90% confluency was reached. Cells were then inoculated with mutant and WT APMV at a MOI of 5. At 1 hpi the inoculum was removed and cells were washed three times with PYG. At 4 or 8 hpi, 10 mL of icecold amoeba salt solution supplemented with 9 g L^−1^ of glucose and 100 µg mL^−1^ of cycloheximide was added to each flask. Cells were then harvested by scraping, pelleted by low-speed centrifugation, washed once in amoeba salt solution, and flash-frozen in LiNi. Cell pellets were lysed on ice using lysis buffer (20 mM Tris-HCl pH 7.4, 100 mM NaCl, 5 mM MgCl_2_, 1% (v/v) Triton X-100, 1 mM DTT, 20 U ml^−1^ RQ1 DNase (Promega, M6101), 0.1% (v/v) NP-40, 100 µg mL^−1^ cycloheximide) and triturated through a 21-gauge needle five times. Preparation of RIBO-seq libraries was carried out by TB-SEQ, Inc. Lysates were digested with RNase I for 45 min at room temp^65^. Digestion was stopped with SuperaseIN and monosomes purified by size exclusion chromatography on MicroSpin S-400 HR columns (GE Healthcare) as described^66^. Size selection of footprints with length 25–33 nt was performed by electrophoresis on 15% TBE-urea gels. Illumina ready RIBO-seq libraries were prepared using a SMARTer smRNA-seq kit (TakaraBio). Library concentrations were measured by qubit fluorometer and their quality assessed on an Agilent 2100 bioanalyzer. RIBO-seq libraries were sequenced on an Illumina Novaseq 6000 sequencer, single read, 1×50 cycles. Preparation of matching RNA-seq libraries was carried out by TB-SEQ, Inc. Total RNA was purified from an aliquot of cell lysate and polyA mRNA prepared using a NEBNext polyA mRNA magnetic isolation module following manufacturer instructions. mRNA fragmentation was conducted for 20 min at 94°C to generate RNA fragments of sizes similar to those of the ribosome footprints. SMARTer smRNA-seq kit (TakaraBio) was used to generate Illumina ready RNA-seq libraries. Library concentrations were measured by qubit fluorometer and their quality assessed on an Agilent 2100 bioanalyzer. RNA-seq libraries were sequenced on an Illumina Novaseq 6000 sequencer, single read, 1×50 cycles. cutadapt v4.6 was used to trim read elements added during library construction (5′ GGG-rich trinucleotide, 3′ adapter sequence, and 3′-end poly-A sequence) with the following custom parameters: -a GATCGGAAGAGCACACGTCTGAACTCCAGTCAC, -a AAAAAAAAAA, -u 3, -m 15. Trimmed reads were then aligned to *A. castellanii* rRNA sequences using STAR v2.7.9a with the following custom parameters: -- outFilterMismatchNoverLmax 0.1, --outFilterMatchNminOverLread 0.9, -- outFilterMatchNmin 14. Reads not mapping to rRNA sequences were then aligned to the APMV genome (NC_014649) and *A. castellanii* nuclear (NW_004457733.1) and mitochondrial genomes (NC_001637.1) accounting for intron/exon gene-structure annotations, with the following custom parameters: --outFilterMatchNmin 15, --outFilterMismatchNoverLmax 0.05, --outFilterMatchNminOverLread 0.95. Read counts were computed using custom scripts to count reads mapped to genomic regions corresponding to annotated rRNA genes, tRNA genes, and to CDS regions of protein-coding genes. After removing mitochondrially encoded genes, samples were normalized for library size and gene length by calculating the transcripts per million (TPM) for each gene. From the sum of TPM values the fraction of reads corresponding to host or viral mRNAs was calculated.

Trimmed reads are available on the GEO database under accession numbers GSE290007. The viral genes highlighted in Fig. 3 and Fig. S4 were selected from the complete RIBO-seq dataset by manually filtering for highly expressed and translated mRNAs with robust annotations, known roles in viral replication, and late expression profiles that correlate with the switch to vIF4F-dependent translation15. Translation efficiencies for these genes were calculated as the ratio of normalized RIBO-seq and RNA-seq TPM for each gene.

### Transmission electron microscopy

*A. castellanii* grown to ∼90% confluency in 150 cm^2^ flasks were infected with APMV WT, *Δ4A, Δ4E*, or *Δ4G* at an MOI of 3. At 1 hpi the inoculum was aspirated and cells were washed three times with PYG. At 12 hpi PYG was replaced with 10 mL ice-cold amoeba salts solution supplemented with 9 g L^−1^ glucose and cells were gently detached by brief exposure to −20°C. Detached cells were pelleted by low-speed centrifugation, washed once with amoeba salts solution, and then fixed overnight at 4°C in a solution containing 2.5% (v/v) glutaraldehyde, 1.25% (v/v) paraformaldehyde and 0.03% (v/v) picric acid in a 0.1 M sodium cacodylate buffer (pH 7.4). Cell pellets were then washed once in 0.1 M sodium cacodylate buffer and postfixed with 1% (v/v) osmium tetroxide and 1.5% (v/v) potassium ferrocyanide for 1 h. Fixed cells were washed in water twice, maleate buffer once, and then incubated in 1% uranyl acetate in maleate buffer for 1 h followed by two washes in water and subsequent dehydration in increasing grades of alcohol (50, 70, 90, and twice at 100%) for 10 min each. The samples were then put in propylene oxide for 1 h and infiltrated overnight in a 1:1 mixture of propylene oxide and TAAB Epon (TAAB Laboratories Equipment Ltd, Aldermaston, England). The following day the samples were embedded in fresh TAAB Epon and polymerized at 60°C for 48 h. Ultrathin sections (∼80 nm) were cut on a Reichert Ultracut-S microtome, picked up on copper grids stained with lead citrate and examined in a JEOL 1200EX Transmission electron microscope equipped with an AMT 2k CCD camera.

### Polysome qRT-PCR

Polysome profiles were generated as described above and ∼1 mL sucrose fractions were collected by hand. Each fraction was diluted 1:1 with RNase-free water before isolating total RNA using phenol-chloroform extraction followed by ethanol precipitation. Briefly, 500 µl of acid phenol-chloroform was added per 500 µl of sucrose solution before vortexing and centrifugation at 21,000 × g for 10 min at 4°C. The aqueous phase was transferred to a new tube containing 50 µl 3M NaOAc and 2 µl glycogen (20 mg mL^−1^) before precipitation at –20°C overnight in 1 mL 100% EtOH. RNA was pelleted by centrifugation at 21,000 × g for 30 min at 4°C. RNA pellets were then washed 3 times with 70% EtOH before resuspending in 14 µL RNase-free water. The entire amount of RNA from each fraction was then used as input for cDNA synthesis using MMLV M5 reverse transcriptase with random hexamer primers. All cDNA reactions were diluted 1:2 with RNase-free water and used as input for qPCR reactions with 2X Luna master mix (NEB, M3003L) and primer sets specific to the viral genes L410, L425, R877 and a validated primer pair targeting the host housekeeping gene HPRT^67^. From the Cq values across each gradient the % RNA of each fraction was calculated as previously described^68^.

### 5′UTR sequence logos

Annotated 5′ UTR sequences from mimivirus (HQ336222.2) and *A. castellanii* transcriptomes (KB007791) were collated, filtered, and trimmed to 10 nt length and used as inputs for WebLogo3^69^.

### Differential scanning fluorimetry

To introduce point mutations, primers with overhangs containing mutant codons were designed. Fragments were generated by PCR and cloned by Gibson assembly into a custom pET16 vector that drives inducible expression of a 6×His-SUMO2 fusion protein. Protein expression and purification was then performed as described above. 20 µl reactions containing 15 µM purified protein, 500 µM ligand, and 3× SYPRO orange dye (ThermoFisher, S6650) were combined on ice in a 96-well plate (ThermoFisher, AB0800WL). All dilutions were made in a buffer containing 20 mM HEPES-KOH pH 7.5 and 250 mM KCl. Samples were then heated from 20°C to 95°C in 0.5°C increments for 30 s using a Real-Time PCR machine (BioRad CFX Opus 96). Fluorescence was measured using SYBR/FAM only scan mode at each increment to generate a melt curve. Melting temperatures were determined using the dRFU tool provided through DSF world^70^.

### Structural AlphaFold-based clustering

Protein clusters generated for the known virome^71^ were used to compare and classify predicted structural features of APMV and Marseilleviridae proteomes. The proteomes of APMV, Marseillevirus, Melbournvirus, and NMV were structurally modeled using ColabFold v1.5.2, executed on the IDRIS Jean Zay GPU supercomputing facility. Multiple sequence alignments were generated on the PACA Bioinfo computing platform using MMseqs2^72^.

### Induction of cell stress

Replication of APMV WT, *Δ4A, Δ4E, Δ4G* and NMV was assessed in *A. castellanii* grown to ∼90% confluency in 6-well plates. To induce ER-stress, cells were treated with 600 nM of thapsigargin (Sigma Aldrich, 67526-95-8) starting 2 h prior to infection or treated with an equal volume of DMSO as control. To induce oxidative stress cells were treated with 50 µM of menadione (Sigma Aldrich, M5625) starting at 1 hpi or treated with an equal volume of DMSO as control. To induce starvation PYG was replaced with amoeba salt solution 22–24 h prior to infection, or left in replete PYG as control. To induce cold shock, cells were transferred to 18°C at 1 hpi or left at 28°C as control. All infections were done at a MOI of 3 and the inoculum was aspirated at 1 hpi followed by three washes with PYG or amoeba salts. All stresses were maintained throughout the experiment until virus-containing supernatants were harvested at 24 hpi and subsequently titered by TCID_50_ as described above. To generate samples for polysome profiles, stressed cells were infected with WT APMV as described above or mock-infected. Acute ER stress was induced through thapsigargin treatment at 7 hpi and cells were then harvested at 8 hpi as described above. Acute starvation was induced by replacement of PYG with amoeba salts at 6 hpi and cells were then harvested at 8 hpi as described above. Polysome profiles were generated and analyzed as described above.

### Quantification and statistical analysis

All information related to the number of biological replicates, definition of center, and dispersion and precision measurements are available in the corresponding figure legend of each figure. All software used for analysis of data is available in the methods details section related to each experiment.

---

**Figure S1.**
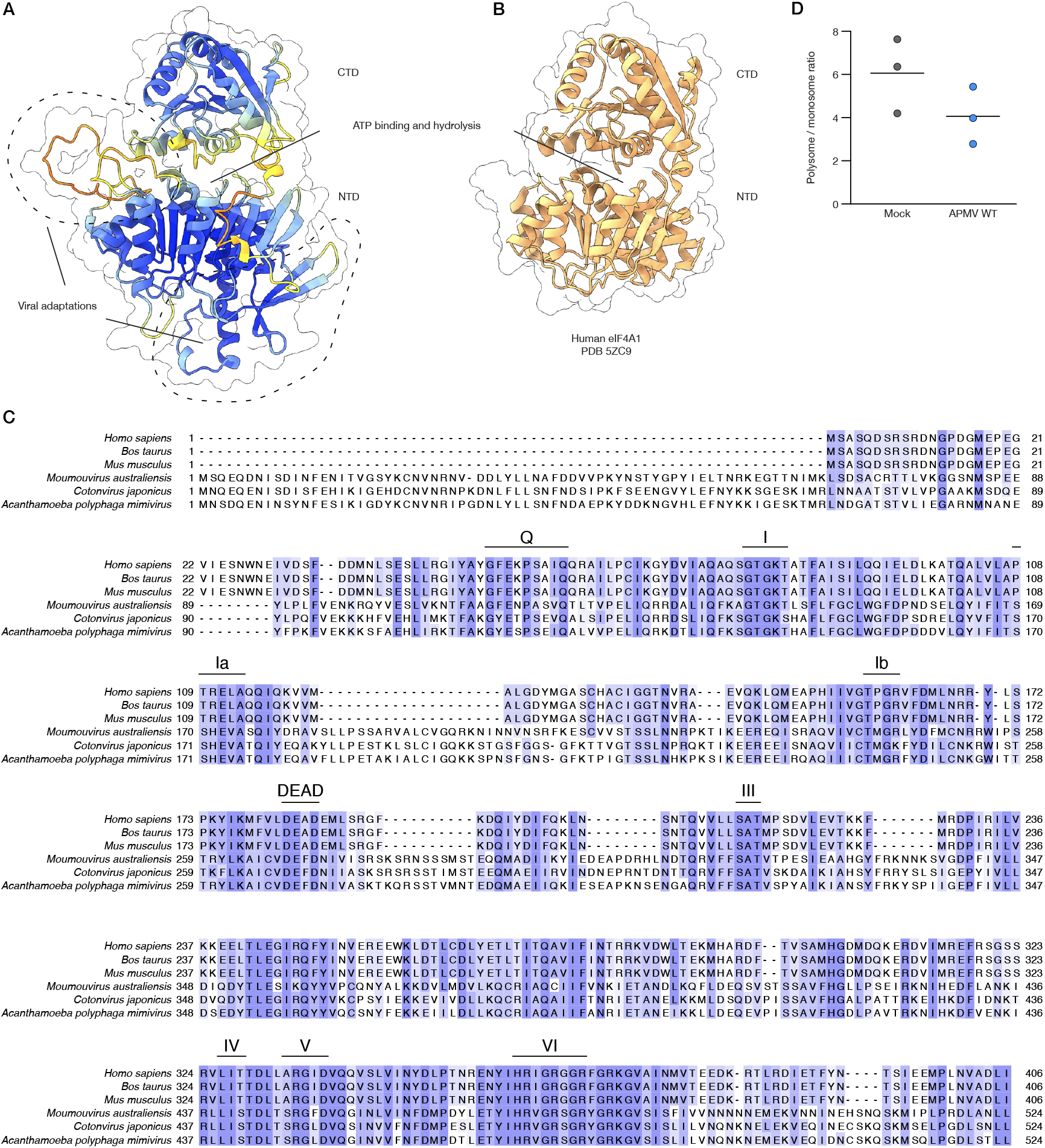
vIF4A is DEAD-box helicase with virus-specific features. **A**. AlphaFold2 model of APMV vIF4A colored by pLDDT. Virus-specific extensions are highlighted. **B**. Crystal structure of human eIF4A1(PDB 5ZC9) illustrating the conserved nature of the two-domain helicase fold. **C**. Multiple sequence alignment of vIF4A and eIF4A colored by % identity. The functional motifs of DEAD-box helicases are highlighted. **D**. Polysome / monosome ratios for mock-infected and APMV WT-infected cells. The line represents the mean of three biological replicates. Related to Figure 1.

**Figure S2.**
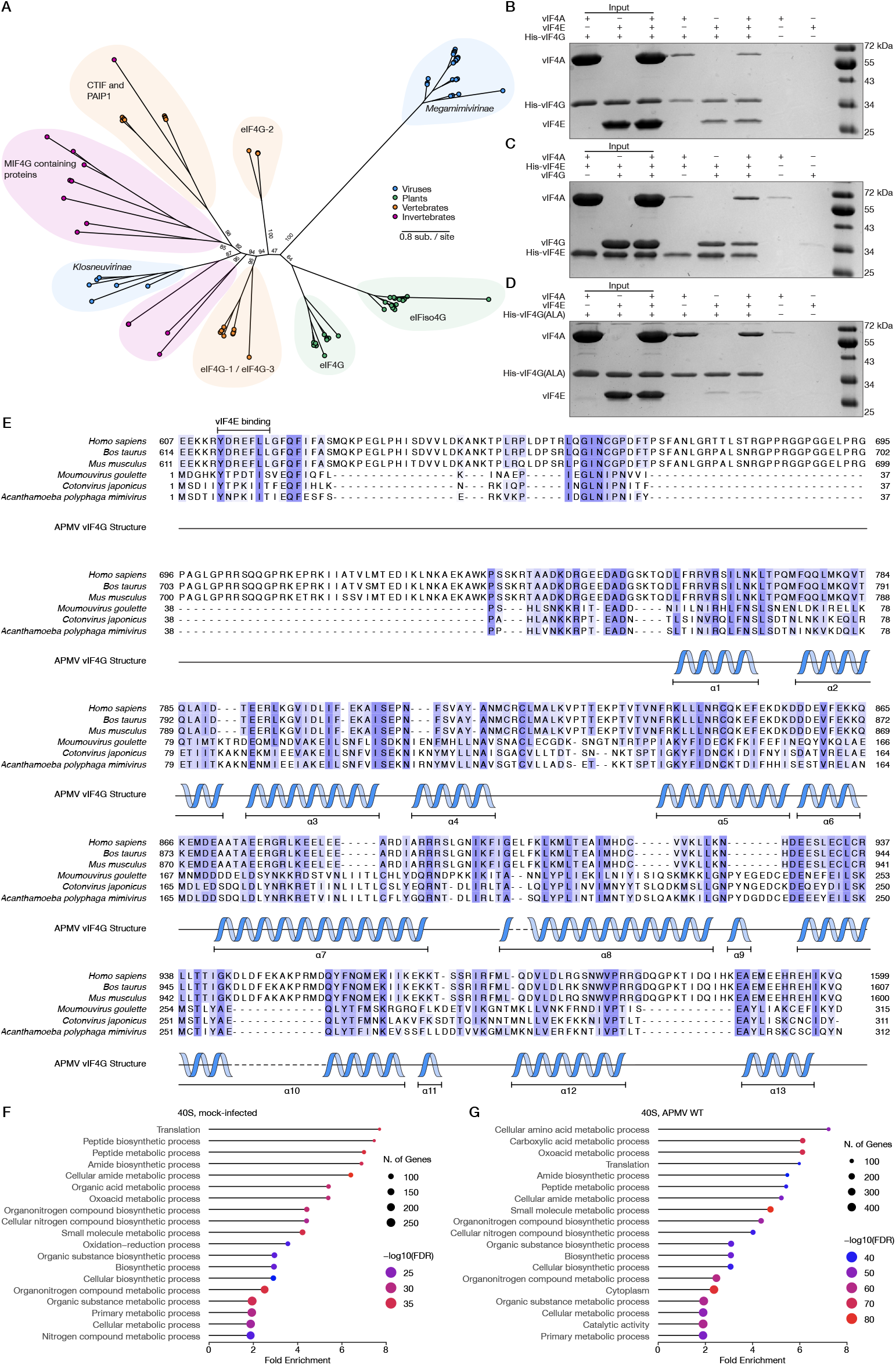
Formation of a viral IF4F complex by a phylogenetically distinct vIF4G. **A**. Phylogenetic tree of eukaryotic and viral MIF4G-like proteins illustrating the divergent nature of APMV vIF4G. Branch supports are provided for the basal branches. **B**. Expanded version of Fig. 1F that includes input lanes and no-bait controls. **C**. vIF4F complex formation mediated by His-vIF4E as visualized by Coomassie-stained SDS-PAGE gel. **D**. Expanded cropped version of Fig. 1G that includes input lanes and no-bait controls. **E**. Multiple sequence alignment of the MIF4G domains from eIF4G1 and vIF4G homologs colored by % identity. The main structural features of APMV vIF4G are aligned with the sequence. **F**. GO enrichment analysis (biological process) of host proteins associated with 40S fractions in mock-infected cells. **G**. GO enrichment analysis (biological process) of host proteins associated with 40S fractions in cells infected by APMV WT. Related to Figure 1.

**Figure S3.**
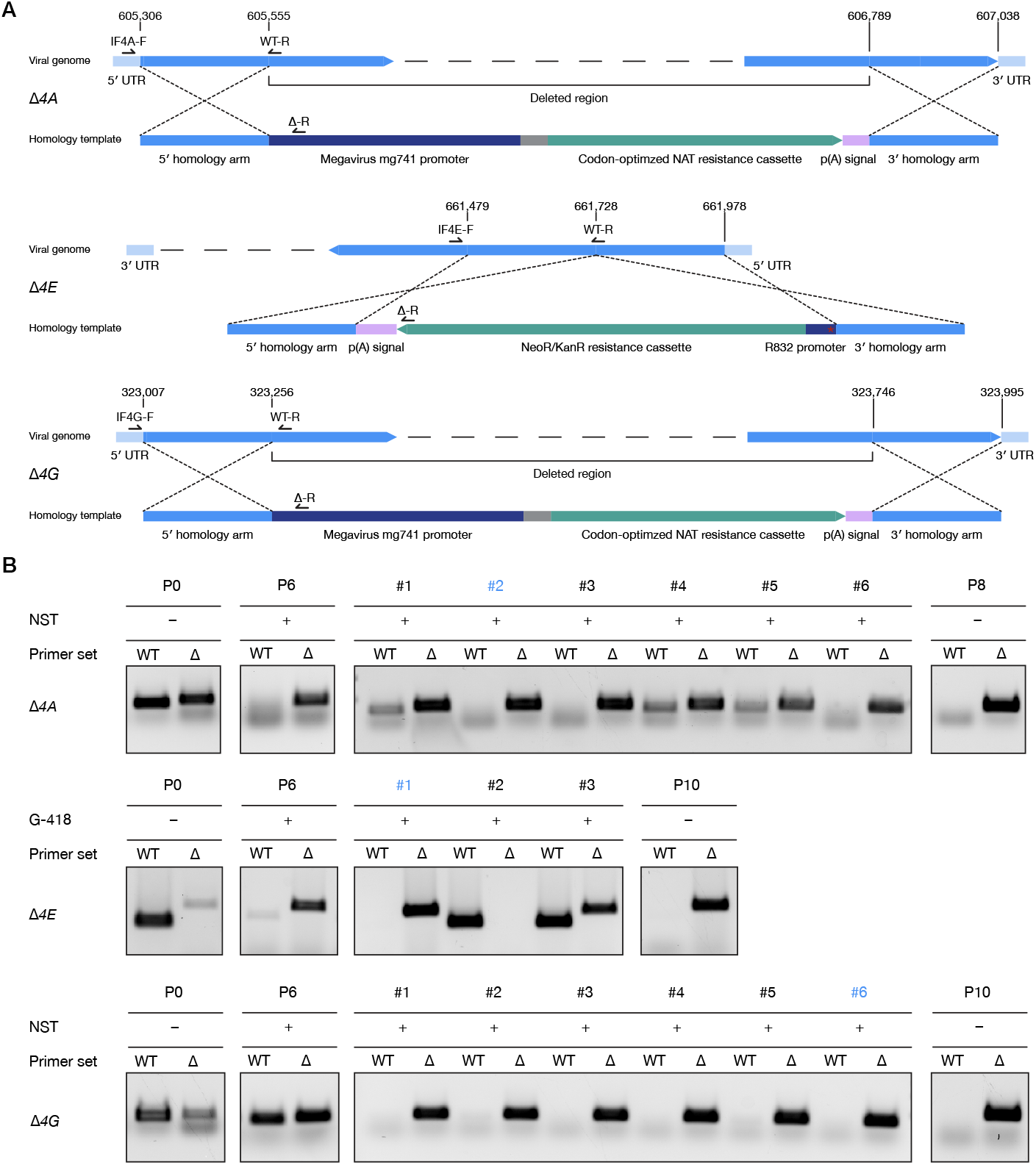
Knock-out of APMV IF4F subunits. **A**. Design of constructs used to generate *Δ4A, Δ4E*, and *Δ4G* APMV. The red star in the R832 promoter indicates a premature stop codon. **B**. Diagnostic PCR of *Δ4A, Δ4E*, and *Δ4G* APMV. The clonal populations selected after limiting dilution are indicated in blue. Related to Figure 2.

**Figure S4.**
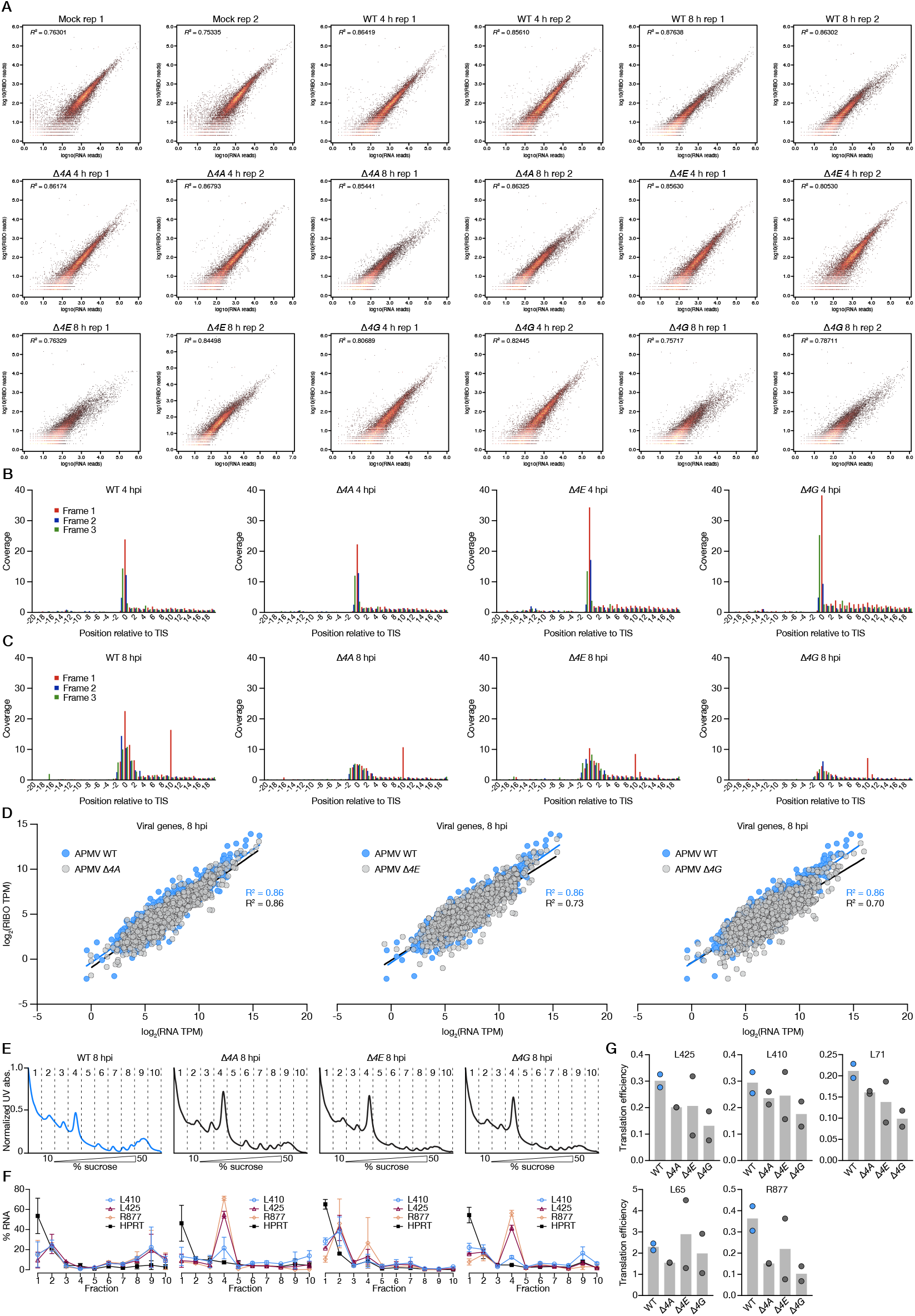
Ribosome profiling quality control and metagene analysis. **A**. Congruence plots of log_10_-transformed RNA-seq and RIBO-seq transcript abundances across samples. Individual replicates are plotted separately. **B**. Metagene analysis, centered on the start codon, of viral genes from WT *Δ4A, Δ4E*, and *Δ4G* APMV at 4 hpi. **C**. Metagene analysis, centered on the start codon, of viral genes from WT *Δ4A, Δ4E*, and *Δ4G* APMV at 8 hpi. **D**. Normalized and log_2_-transformed RNA and RPF abundances of all viral transcripts detected at 8 hpi during APMV *Δ4A, Δ4E*, and *Δ4G* infection. Distributions of the mutants are overlaid onto that of APMV WT for visualization purposes. Each dot represents the mean of two biological replicates and solid lines represent a linear fit of the data with the associated R^2^ values. **E**. Representative examples of polysome profiles from cells infected with APMV WT, *Δ4A, Δ4E*, or *Δ4G* collected at 8 hpi. Sucrose gradient fractions are marked by dashed lines. **F**. % of target mRNAs associated with each sucrose gradient fraction from cells infected with APMV WT, *Δ4A, Δ4E*, or *Δ4G* at 8 hpi. Symbols and lines represent the mean +/-range from two biological replicates. **G**. Translation efficiency (RIBO-seq: RNA-seq ratio) for each of the selected structural proteins in cells infected with APMV WT, *Δ4A, Δ4E*, or *Δ4G* at 8 hpi. Each bar represents the mean of two biological replicates. Related to Figure 3.

**Figure S5.**
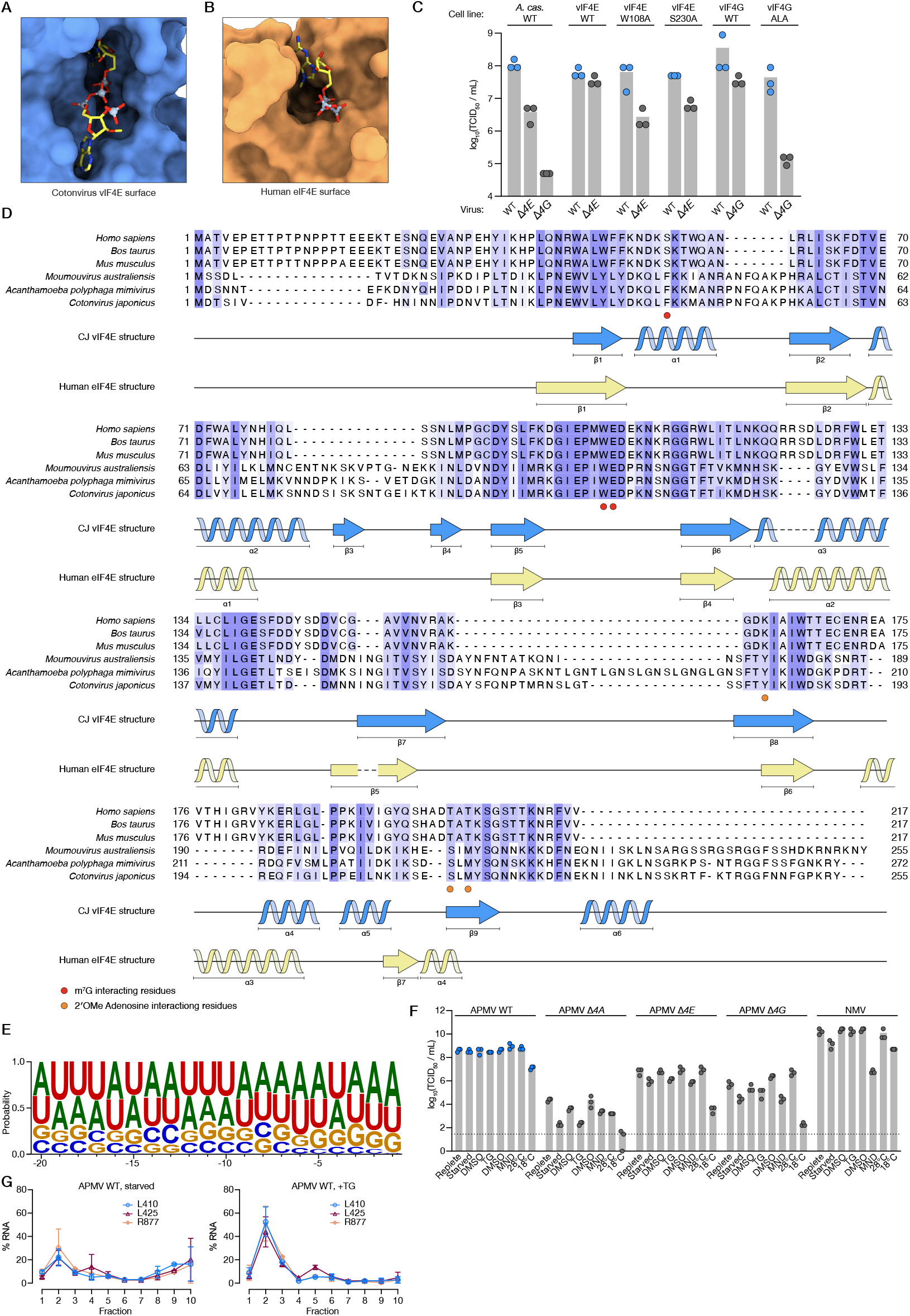
Structural and sequence comparison of vIF4E and human eIF4E. **A**. Surface representation of vIF4E from CJ in complex with m^7^Gppp(2′OMeA)pU demonstrating the close coordination of the ligand. **B**. Surface representation of human eIF4E in complex with m^7^Gppp (PDB 5T46). **C**. Viral replication in *A. castellanii* cell lines expressing mutant or WT forms of vIF4E and vIF4G. Each bar represents the mean of three biological replicates. All residue numbers correspond to APMV vIF4E. **D**. Multiple sequence alignment of the vIF4E and eIF4E homologs colored by % identity. The main structural features of CJ vIF4E and human eIF4E are aligned with the sequence. **E**. Weblogo of the 20 nt upstream of the AUG in NMV intergenic regions. **F**. Raw titers of APMV WT, *Δ4A, Δ4E, Δ4G*, or NMV replication under stress conditions. The limit of detection is indicated by a dotted line. **G**. % of target mRNAs associated with each sucrose gradient fraction from cells infected with APMV WT under stravation or TG treatment 8 hpi. Related to Figure 4 and 5.

## References

1. Bernier, C.R., Petrov, A.S., Kovacs, N.A., Penev, P.I., and Williams, L.D. (2018). Translation: The Universal Structural Core of Life. Mol. Biol. Evol. 35, 2065–2076. 10.1093/molbev/msy101.

2. Bowman, J.C., Petrov, A.S., Frenkel-Pinter, M., Penev, P.I., and Williams, L.D. (2020). Root of the Tree: The Significance, Evolution, and Origins of the Ribosome. Chem. Rev. 120, 4848–4878. 10.1021/acs.chemrev.9b00742.

3. Gross, J.D., Moerke, N.J., Haar, T. von der, Lugovskoy, A.A., Sachs, A.B., McCarthy, J.E.G., and Wagner, G. (2003). Ribosome Loading onto the mRNA Cap Is Driven by Conformational Coupling between eIF4G and eIF4E. Cell 115, 739–750. 10.1016/S0092-8674(03)00975-9.

4. Linder, P., Lasko, P.F., Ashburner, M., Leroy, P., Nielsen, P.J., Nishi, K., Schnier, J., and Slonimski, P.P. (1989). Birth of the D-E-A-D box. Nature 337, 121–122. 10.1038/337121a0.

5. Sonenberg, N., Rupprecht, K.M., Hecht, S.M., and Shatkin, A.J. (1979). Eukaryotic mRNA cap binding protein: purification by affinity chromatography on sepharose-coupled m7GDP. Proc. Natl. Acad. Sci. U. S. A. 76, 4345–4349.

6. Marintchev, A., Edmonds, K.A., Marintcheva, B., Hendrickson, E., Oberer, M., Suzuki, C., Herdy, B., Sonenberg, N., and Wagner, G. (2009). Topology and Regulation of the Human eIF4A/4G/4H Hel-icase Complex in Translation Initiation. Cell 136, 447–460. 10.1016/j.cell.2009.01.014.

7. Lwoff, A. (1957). The Concept of Virus. Microbiology 17, 239–253. 10.1099/00221287-17-2-239.

8. Scola, B.L., Audic, S., Robert, C., Jungang, L., de Lamballerie, X., Drancourt, M., Birtles, R., Claverie, J.-M., and Raoult, D. (2003). A Giant Virus in Amoebae. Science 299, 2033–2033. 10.1126/science.1081867.

9. Raoult, D., Audic, S., Robert, C., Abergel, C., Renesto, P., Ogata, H., La Scola, B., Suzan, M., and Claverie, J.-M. (2004). The 1.2-Megabase Genome Sequence of Mimivirus. Science 306, 1344–1350. 10.1126/science.1101485.

10. Schulz, F., Yutin, N., Ivanova, N.N., Ortega, D.R., Lee, T.K., Vierheilig, J., Daims, H., Horn, M., Wagner, M., Jensen, G.J., et al. (2017). Giant viruses with an expanded complement of translation system components. Science 356, 82–85. 10.1126/science.aal4657.

11. Abrahão, J., Silva, L., Silva, L.S., Khalil, J.Y.B., Rodrigues, R., Arantes, T., Assis, F., Boratto, P., Andrade, M., Kroon, E.G., et al. (2018). Tailed giant Tupanvirus possesses the most complete translational apparatus of the known virosphere. Nat. Commun. 9, 749. 10.1038/s41467-018-03168-1.

12. Simsek, D., Tiu, G.C., Flynn, R.A., Byeon, G.W., Leppek, K., Xu, A.F., Chang, H.Y., and Barna, M. (2017). The Mammalian Ribo-interactome Reveals Ribosome Functional Diversity and Heterogeneity. Cell 169, 1051-1065.e18. 10.1016/j.cell.2017.05.022.

13. Takahashi, H., Fukaya, S., Song, C., Murata, K., and Takemura, M. (2021). Morphological and Taxonomic Properties of the Newly Isolated Cotonvirus japonicus, a New Lineage of the Subfamily Megavirinae. J. Virol. 95, 10.1128/jvi.00919-21. 10.1128/jvi.00919-21.

14. Mader, S., Lee, H., Pause, A., and Sonenberg, N. (1995). The translation initiation factor eIF-4E binds to a common motif shared by the translation factor eIF-4 gamma and the translational repressors 4Ebinding proteins. Mol. Cell. Biol. 15, 4990–4997.

15. Legendre, M., Audic, S., Poirot, O., Hingamp, P., Seltzer, V., Byrne, D., Lartigue, A., Lescot, M., Bernadac, A., Poulain, J., et al. (2010). mRNA deep sequencing reveals 75 new genes and a complex transcriptional landscape in Mimivirus. Genome Res. 20, 664–674. 10.1101/gr.102582.109.

16. Marcotrigiano, J., Gingras, A.-C., Sonenberg, N., and Burley, S.K. (1997). Cocrystal Structure of the Messenger RNA 5′ Cap-Binding Protein (eIF4E) Bound to 7-methyl-GDP. Cell 89, 951 – 961. 10.1016/S0092-8674(00)80280-9.

17. Dix, T.C., Haussmann, I.U., Brivio, S., Nallasivan, M.P., HadzHiev, Y., Müller, F., Müller, B., Pettitt, J., and Soller, M. (2022). CMTr mediated 2′-O-ribose methylation status of cap-adjacent nucleotides across animals. RNA 28, 1377–1390. 10.1261/rna.079317.122.

18. Benarroch, D., Smith, P., and Shuman, S. (2008). Characterization of a trifunctional mimivirus mRNA capping enzyme and crystal structure of the RNA triphosphatase domain. Struct. Lond. Engl. 1993 16, 501– 512. 10.1016/j.str.2008.01.009.

19. Priet, S., Lartigue, A., Debart, F., Claverie, J.-M., and Abergel, C. (2015). mRNA maturation in giant viruses: variation on a theme. Nucleic Acids Res. 43, 3776–3788. 10.1093/nar/gkv224.

20. Daffis, S., Szretter, K.J., Schriewer, J., Li, J., Youn, S., Errett, J., Lin, T.-Y., Schneller, S., Zust, R., Dong, H., et al. (2010). 2′-O methylation of the viral mRNA cap evades host restriction by IFIT family members. Nature 468, 452–456. 10.1038/nature09489.

21. Devarkar, S.C., Wang, C., Miller, M.T., Ramanathan, A., Jiang, F., Khan, A.G., Patel, S.S., and Marcotrigiano, J. (2016). Structural basis for m7G recognition and 2’-O-methyl discrimination in capped RNAs by the innate immune receptor RIG-I. Proc. Natl. Acad. Sci. U. S. A. 113, 596–601. 10.1073/pnas.1515152113.

22. Picard-Jean, F., Brand, C., Tremblay-Létourneau, M., Allaire, A., Beaudoin, M.C., Boudreault, S., Duval, C., Rainville-Sirois, J., Robert, F., Pelletier, J., et al. (2018). 2’-O-methylation of the mRNA cap protects RNAs from decapping and degradation by DXO. PloS One 13, e0193804. 10.1371/journal.pone.0193804.

23. Grüner, S., Peter, D., Weber, R., Wohlbold, L., Chung, M.-Y., Weichenrieder, O., Valkov, E., Igreja, C., and Izaurralde, E. (2016). The Structures of eIF4E-eIF4G Complexes Reveal an Extended Interface to Regulate Translation Initiation. Mol. Cell 64, 467–479. 10.1016/j.molcel.2016.09.020.

24. Frydryskova, K., Masek, T., Borcin, K., Mrvova, S., Venturi, V., and Pospisek, M. (2016). Distinct recruitment of human eIF4E isoforms to processing bodies and stress granules. BMC Mol. Biol. 17, 21. 10.1186/s12867-016-0072-x.

25. Ptushkina, M., Malys, N., and McCarthy, J.E.G. (2004). eIF4E isoform 2 in Schizosaccharomyces pombe is a novel stress-response factor. EMBO Rep. 5, 311–316. 10.1038/sj.embor.7400088.

26. Felipe Benites, L., Stephens, T.G., Van Etten, J., James, T., Christian, W.C., Barry, K., Grigoriev, I.V., McDermott, T.R., and Bhattacharya, D. (2024). Hot springs viruses at Yellowstone National Park have ancient origins and are adapted to thermophilic hosts. Commun. Biol. 7, 1–12. 10.1038/s42003-024-05931-1.

27. Rigou, S., Santini, S., Abergel, C., Claverie, J.-M., and Legendre, M. (2022). Past and present giant viruses diversity explored through permafrost metagenomics. Nat. Commun. 13, 5853. 10.1038/s41467-022-33633-x.

28. McGarry, T.J., and Lindquist, S. (1985). The preferential translation of Drosophila hsp70 mRNA requires sequences in the untranslated leader. Cell 42, 903–911. 10.1016/0092-8674(85)90286-7.

29. Lee, A.S.Y., Kranzusch, P.J., Doudna, J.A., and Cate, J.H.D. (2016). eIF3d is an mRNA cap-binding protein that is required for specialized translation initiation. Nature 536, 96–99. 10.1038/nature18954.

30. Mukhopadhyay, S., Amodeo, M.E., and Lee, A.S.Y. (2023). eIF3d controls the persistent integrated stress response. Mol. Cell 83, 33033313.e6. 10.1016/j.molcel.2023.08.008.

31. Lamper, A.M., Fleming, R.H., Ladd, K.M., and Lee, A.S.Y. (2020). A phosphorylation-regulated eIF3d translation switch mediates cellular adaptation to metabolic stress. Science 370, 853–856. 10.1126/science.abb0993.

32. Pause, A., Belsham, G.J., Gingras, A.C., Donzé, O., Lin, T.A., Lawrence, J.C., and Sonenberg, N. (1994). Insulin-dependent stimulation of protein synthesis by phosphorylation of a regulator of 5’-cap function. Nature 371, 762–767. 10.1038/371762a0.

33. Aylward, F.O., Moniruzzaman, M., Ha, A.D., and Koonin, E.V. (2021). A phylogenomic framework for charting the diversity and evolution of giant viruses. PLOS Biol. 19, e3001430. 10.1371/journal.pbio.3001430.

34. Fabre, E., Jeudy, S., Santini, S., Legendre, M., Trauchessec, M., Couté, Y., Claverie, J.-M., and Abergel, C. (2017). Noumeavirus replication relies on a transient remote control of the host nucleus. Nat. Commun. 8, 15087. 10.1038/ncomms15087.

35. Oliveira, G.P., Lima, M.T., Arantes, T.S., Assis, F.L., Rodrigues, R.A.L., da Fonseca, F.G., Bonjardim, C.A., Kroon, E.G., Colson, P., La Scola, B., et al. (2017). The Investigation of Promoter Sequences of Marseilleviruses Highlights a Remarkable Abundance of the AAATATTT Motif in Intergenic Regions. J. Virol. 91, 10.1128/jvi.01088-17. 10.1128/jvi.01088-17.

36. Suhre, K., Audic, S., and Claverie, J.-M. (2005). Mimivirus gene promoters exhibit an unprecedented conservation among all eukaryotes. Proc. Natl. Acad. Sci. 102, 14689–14693. 10.1073/pnas.0506465102.

37. Rigou, S., Schmitt, A., Moreno, A.B., Lartigue, A., Danner, L., Giry, C., Trabelsi, F., Belmudes, L., Olivero-Deibe, N., Guenno, H.L., et al. (2025). Nucleocytoviricota viral factories are transient organelles made by liquid-liquid phase separation. bioRxiv. 10.1101/2024.09.01.610734.

38. Neidermyer, W.J., and Whelan, S.P.J. (2019). Global analysis of polysome-associated mRNA in vesicular stomatitis virus infected cells. PLoS Pathog. 15, e1007875. 10.1371/journal.ppat.1007875.

39. Garrey, J.L., Lee, Y.-Y., Au, H.H.T., Bushell, M., and Jan, E. (2010). Host and Viral Translational Mechanisms during Cricket Paralysis Virus Infection. J. Virol. 84, 1124–1138. 10.1128/JVI.02006-09.

40. Aravind, L., and Koonin, E.V. (2000). Eukaryote-specific Domains in Translation Initiation Factors: Implications for Translation Regulation and Evolution of the Translation System. Genome Res. 10, 1172– 1184.

41. Kyrpides, N.C., and Woese, C.R. (1998). Universally conserved translationinitiationfactors. Proc. Natl. Acad. Sci. U. S. A. 95, 224–228.

42. Hernández, G., and Vazquez-Pianzola, P. (2005). Functional diversity of the eukaryotic translation initiation factors belonging to eIF4 families. Mech. Dev. 122, 865–876. 10.1016/j.mod.2005.04.002.

43. Gentry, R.C., Ide, N.A., Comunale, V.M., Hartwick, E.W., Kinz-Thompson, C.D., and Gonzalez, R.L. (2024). The mechanism of mRNA cap recognition. Nature, 1–8. 10.1038/s41586-024-08304-0.

44. Mayer, L., Nikolov, G., Kunert, M., Horn, M., and Willemsen, A. (2025). Giant virus transcription and translation occur at well-defined locations within amoeba host cells. Preprint, 10.1101/2025.03.24.645094 10.1101/2025.03.24.645094.

45. Zhang, R., Mayer, L., Hikida, H., Shichino, Y., Mito, M., Willemsen, A., Iwasaki, S., and Ogata, H. (2024). Giant virus creates subcellular environment to overcome codon– tRNA mismatch. Preprint, 10.1101/2024.10.07.616867 10.1101/2024.10.07.616867.

46. Abrahão, J.S., Boratto, P., Dornas, F.P., Silva, L.C., Campos, R.K., Almeida, G.M.F., Kroon, E.G., and La Scola, B. (2014). Growing a giant: Evaluation of the virological parameters for mimivirus production. J. Virol. Methods 207, 6–11. 10.1016/j.jviromet.2014.06.001.

47. Lei, C., Yang, J., Hu, J., and Sun, X. (2021). On the Calculation of TCID50 for Quantitation of Virus Infectivity. Virol. Sin. 36, 141–144. 10.1007/s12250-020-00230-5.

48. Philippe, N., Shukla, A., Abergel, C., and Bisio, H. (2024). Genetic manipulation of giant viruses and their host, Acanthamoeba castellanii. Nat. Protoc. 19, 3–29. 10.1038/s41596-023-00910-y.

49. Schiller, C. QuAPPro: An R/shiny app for Quantification and Alignment of Polysome Profiles | bioRxiv. https://www.biorxiv.org/con-tent/10.1101/2024.05.02.592260v1.full.

50. Shevchenko, A., Wilm, M., Vorm, O., and Mann, M. (1996). Mass spectrometric sequencing of proteins silver-stained polyacrylamide gels. Anal. Chem. 68, 850–858. 10.1021/ac950914h.

51. Peng, J., and Gygi, S.P. (2001). Proteomics: the move to mixtures. J. Mass Spectrom. JMS 36, 1083–1091. 10.1002/jms.229.

52. Eng, J.K., McCormack, A.L., and Yates, J.R. (1994). An approach to correlate tandem mass spectral data of peptides with amino acid sequences in a protein database. J. Am. Soc. Mass Spectrom. 5, 976–989. 10.1016/1044-0305(94)80016-2.

53. Mirdita, M., Schütze, K., Moriwaki, Y., Heo, L., Ovchinnikov, S., and Steinegger, M. (2022). ColabFold: making protein folding accessible to all. Nat. Methods 19, 679–682. 10.1038/s41592-022-01488-1.

54. van Kempen, M., Kim, S.S., Tumescheit, C., Mirdita, M., Lee, J., Gilchrist, C.L.M., Söding, J., and Steinegger, M. (2024). Fast and accurate protein structure search with Foldseek. Nat. Biotechnol. 42, 243– 246. 10.1038/s41587-023-01773-0.

55. Sievers, F., Wilm, A., Dineen, D., Gibson, T.J., Karplus, K., Li, W., Lopez, R., McWilliam, H., Remmert, M., Söding, J., et al. (2011). Fast, scalable generation of high-quality protein multiple sequence alignments using Clustal Omega. Mol. Syst. Biol. 7, 539. 10.1038/msb.2011.75.

56. Price, M.N., Dehal, P.S., and Arkin, A.P. (2009). FastTree: Computing Large Minimum Evolution Trees with Profiles instead of a Distance Matrix. Mol. Biol. Evol. 26, 1641–1650. 10.1093/molbev/msp077.

57. Price, M.N., Dehal, P.S., and Arkin, A.P. (2010). FastTree 2 – Approximately Maximum-Likelihood Trees for Large Alignments. PLOS ONE 5, e9490. 10.1371/journal.pone.0009490.

58. Waterhouse, A.M., Procter, J.B., Martin, D.M.A., Clamp, M., and Barton, G.J. (2009). Jalview Version 2--a multiple sequence alignment editor and analysis workbench. Bioinforma. Oxf. Engl. 25, 1189–1191. 10.1093/bioinformatics/btp033.

59. Ge, S.X., Jung, D., and Yao, R. (2020). ShinyGO: a graphical gene-set enrichment tool for animals and plants. Bioinforma. Oxf. Engl. 36, 2628–2629. 10.1093/bioinformatics/btz931.

60. Zhou, W., Whiteley, A.T., de Oliveira Mann, C.C., Morehouse, B.R., Nowak, R.P., Fischer, E.S., Gray, N.S., Mekalanos, J.J., and Kranzusch, P.J. (2018). Structure of the Human cGAS-DNA Complex Reveals Enhanced Control of Immune Surveillance. Cell 174, 300-311.e11. 10.1016/j.cell.2018.06.026.

61. Vonrhein, C., Flensburg, C., Keller, P., Sharff, A., Smart, O., Paciorek, W., Womack, T., and Bricogne, G. (2011). Data processing and analysis with the autoPROC toolbox. Acta Crystallogr. D Biol. Crystallogr. 67, 293–302. 10.1107/S0907444911007773.

62. Emsley, P., Lohkamp, B., Scott, W.G., and Cowtan, K. (2010). Features and development of Coot. Acta Crystallogr. D Biol. Crystallogr. 66, 486–501. 10.1107/S0907444910007493.

63. Liebschner, D., Afonine, P.V., Baker, M.L., Bunkóczi, G., Chen, V.B., Croll, T.I., Hintze, B., Hung, L.-W., Jain, S., McCoy, A.J., et al. (2019). Macromolecular structure determination using X-rays, neutrons and electrons: recent developments in Phenix. Acta Crystallogr. Sect. Struct. Biol. 75, 861–877. 10.1107/S2059798319011471.

64. Meng, E.C., Goddard, T.D., Pettersen, E.F., Couch, G.S., Pearson, Z.J., Morris, J.H., and Ferrin, T.E. (2023). UCSF ChimeraX: Tools for structure building and analysis. Protein Sci. Publ. Protein Soc. 32, e4792. 10.1002/pro.4792.

65. McGlincy, N.J., and Ingolia, N.T. (2017). Transcriptome-wide measurement of translation by ribosome profiling. Methods San Diego Calif 126, 112–129. 10.1016/j.ymeth.2017.05.028.

66. Aeschimann, F., Xiong, J., Arnold, A., Dieterich, C., and Großhans, H. (2015). Transcriptome-wide measurement of ribosomal occupancy by ribosome profiling. Methods San Diego Calif 85, 75–89. 10.1016/j.ymeth.2015.06.013.

67. Köhsler, M., Leitsch, D., Müller, N., and Walochnik, J. (2020). Validation of reference genes for the normalization of RT-qPCR gene expression in Acanthamoeba spp. Sci. Rep. 10, 10362. 10.1038/s41598-020-67035-0.

68. Han, C., Sun, L., Pan, Q., Sun, Y., Wang, W., and Chen, Y. (2021). Polysome profiling followed by quantitative PCR for identifying potential micropeptide encoding long non-coding RNAs in suspension cell lines. STAR Protoc. 3, 101037. 10.1016/j.xpro.2021.101037.

69. Crooks, G.E., Hon, G., Chandonia, J.-M., and Brenner, S.E. (2004). WebLogo: a sequence logo generator. Genome Res. 14, 1188–1190. 10.1101/gr.849004.

70. Wu, T., Gale-Day, Z.J., and Gestwicki, J.E. (2024). DSFworld: A flexible and precise tool to analyze differential scanning fluorimetry data. Protein Sci. Publ. Protein Soc. 33, e5022. 10.1002/pro.5022.

71. Nomburg, J., Doherty, E.E., Price, N., Bellieny-Rabelo, D., Zhu, Y.K., and Doudna, J.A. (2024). Birth of protein folds and functions in the virome. Nature 633, 710–717. 10.1038/s41586-024-07809-y.

72. Steinegger, M., and Söding, J. (2017). MMseqs2 enables sensitive protein sequence searching for the analysis of massive data sets. Nat. Biotechnol. 35, 1026–1028. 10.1038/nbt.3988.

